# Intracellular pH differentially regulates transcription of metabolic and signaling pathways in normal epithelial cells

**DOI:** 10.1101/2022.07.12.499804

**Authors:** Ricardo Romero-Moreno, Brandon Czowski, Lindsey Harris, Jessamine F. Kuehn, Katharine A. White

## Abstract

Intracellular pH (pHi) dynamics regulate normal cell function, and dysregulated pHi dynamics is an emerging hallmark of cancer (constitutively increased pHi) and neurodegeneration (constitutively decreased pHi). However, the molecular mechanisms by which pHi dynamics regulate cell biology are poorly understood. Here, we discovered that altering pHi in normal human breast epithelial cells triggers global transcriptional changes. We identified 176 genes differentially regulated by pHi, with pHi-dependent genes clustering in signaling and glycolytic pathways. Using various normal epithelial cell models, we showed pH-dependent Notch1 expression, with increased protein abundance at high pHi. This resulted in pH-dependent downstream signaling, with increased Notch1 signaling at high pHi. We also found that high pHi increased the expression of glycolytic enzymes and regulators of pyruvate fate, including lactate dehydrogenase and pyruvate dehydrogenase kinase. These transcriptional changes were sufficient to alter lactate production, with high pHi shifting these normal epithelial cells toward a glycolytic metabolism and increasing lactate production. Thus, pHi dynamics transcriptionally regulate signaling and metabolic pathways in normal epithelial cells. Our data reveal new molecular regulators of pHi-dependent biology and a role for increased pHi in driving the acquisition of cancer-associated signaling and metabolic changes in normal human epithelial cells.

## Introduction

Normal epithelial cells have an intracellular pH (pHi) of ∼7.2, but transient changes in pHi (6.8-7.6) have been shown to regulate cell behaviors, including cell migration (1), proliferation (2, 3), and differentiation (4) (Figure 1A). Transient pHi dynamics are driven by regulation of the expression and activity of ion transport proteins, including monocarboxylate transporters(5), sodium bicarbonate transporters (6), V-ATPases (7), the Na^+^-H^+^ exchanger 1 (NHE1) (8), and anion exchangers (9, 10). At the molecular scale, pH-dependent biology is mediated by proteins or pathways that are sensitive to small, physiological increases (7.2-7.6) or decreases (6.8-7.2) in pHi (11). The subclass of proteins with activities, abundances, or binding affinities that are sensitive to physiological changes in pHi are called pH sensors.

**Figure 1.**
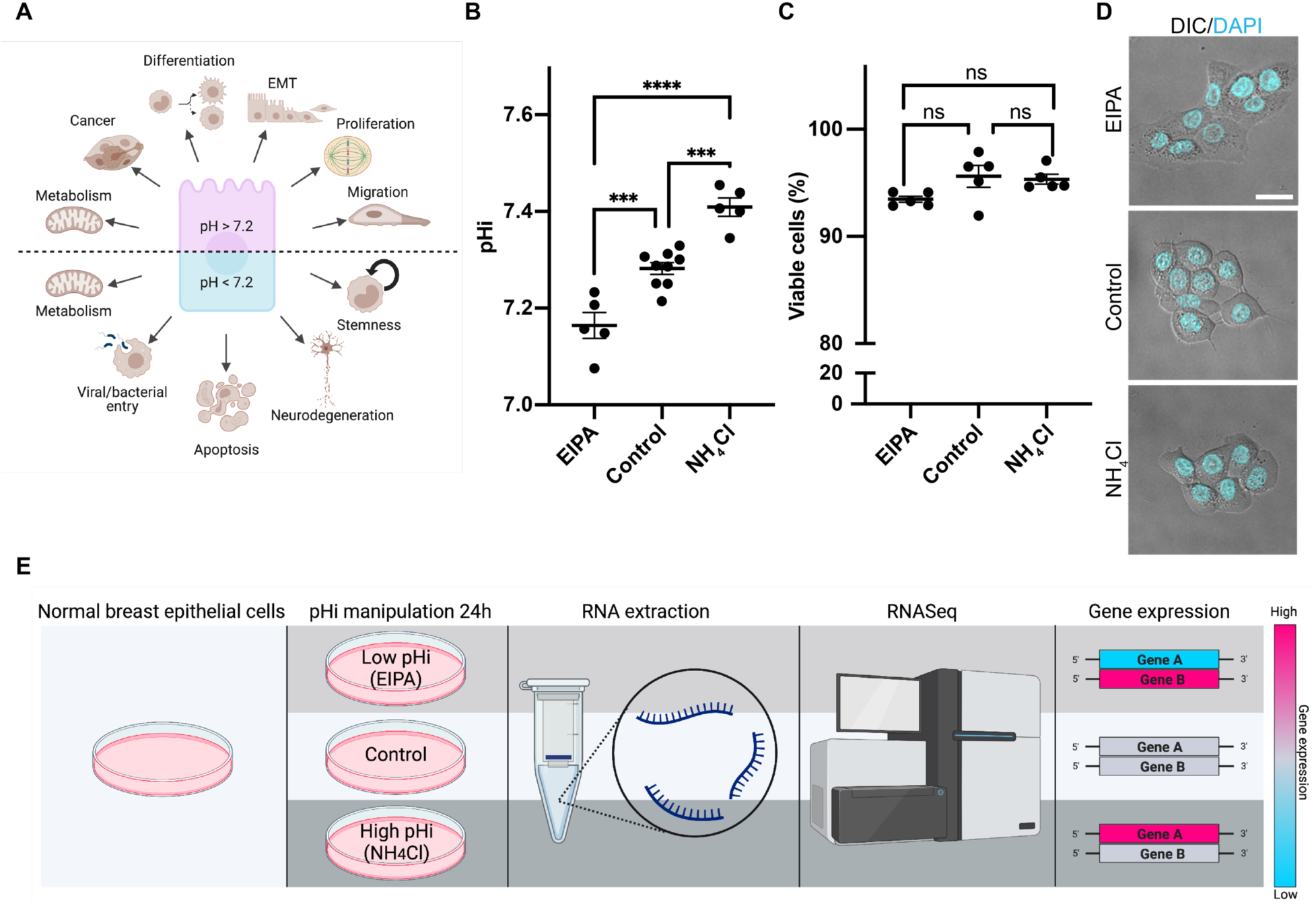
Increasing and decreasing pHi in normal breast epithelial cells does not alter viability or morphology. A) Schematic showing pHi-dependent biology and dysregulated pHi of cancer and neurodegeneration. B) Measured pHi (see methods) in normal human mammary epithelial cells (MCF10A) that were left untreated (control) or treated for 24 hours with complete media containing 25 μM EIPA (EIPA) to lower pHi or 30 mM NH4Cl (NH4Cl) to raise pHi. Scatter plot (mean±SEM) from 5-9 biological replicates. C) Trypan blue exclusion assay to assess the viability of cells treated as in B. Scatter plot (mean±SEM) from 5 biological replicates. D) Representative confocal images of MCF10A cells treated as in (B). Differential interference contrast (DIC), with overlay of Hoechst dye nuclear stain (DAPI). Scale bars: 20 μm. E) Schematic of pHi manipulation, RNA isolation, and analysis workflow. Resulting RNASeq analysis can identify genes that are upregulated (magenta) or downregulated (cyan) compared to control MCF10A cells. For B and C, significance was determined using ANOVA with Tukey’s multiple comparisons correction, ***p<0.001, ****p<0.0001.

Work identifying pH sensors and pH-sensitive pathways has generally started from the pH-dependent biological outcome, identified essential pathways in that process, and then tested the pH sensitivity of critical pathway proteins one by one. This laborious approach has demonstrated successes with the identification of actin cytoskeleton remodeling as a pH-sensitive pathway in Dictyostelium in 1996 (12), mammalian cells in 2002 (1), and finally the identification of four pH-sensitive proteins involved in actin cytoskeleton remodeling (talin (13), cofilin (14), cdc42 (15), and focal adhesion kinase (16) with full molecular mechanisms characterized.

However, for many pH-sensitive behaviors, such as cell-cycle progression (2, 3) and epithelial to mesenchymal transition (17), metabolic regulation (18, 19), and differentiation (4) the mediating pH sensors are unknown or uncharacterized (11). One way to more rapidly reveal the molecular mechanisms driving pHi-dependent biology is by applying −omics-based approaches to rapidly identify pH-dependent proteins or pH-sensitive pathway nodes. With the goal of identifying global pH-dependent transcriptional regulation, prior work by Putney *et al.*, prepared a normal mouse fibroblast cell line where a ubiquitously-expressed pH homeostatic regulator, Na^+^-H^+^ exchanger (NHE1), was deleted (20). This cell line was then supplemented with either wild-type NHE1 to produce a cell line (LAPN) with a normal pHi of 7.35 or a transport-dead NHE1 variant (NHE1-E266I) to produce a cell line (LAPE) with a decreased pHi of 7.16 (20). The authors then showed that these two cell models had significantly altered expression profiles of transcripts related to cell cycle regulation, growth, and metabolic adaptation (20).

While this dataset established a role for intracellular pH in transcriptional regulation of normal cells, the work has four significant limitations. First, Putney *et al.* used a mouse fibroblast cell line, which does not replicate normal human epithelial biology. Second, Putney *et al.* only compared low pHi to normal pHi, which is a substantial limitation given that many pH-dependent processes with unknown molecular drivers (e.g., epithelial to mesenchymal transition, cell differentiation, proliferation) (2–4, 21) involve transient increases in pHi. Third, Putney *et al.* used an Affymetrix microarray, which has limited transcriptome coverage compared to modern NextGen RNASeq approaches. Finally, Putney *et al.* used genetic deletion and supplementation to generate the two mouse fibroblast cell lines with normal and low pHi homeostasis. Thus, the results are confounded by off-target compensatory effects arising from NHE1 deletion followed by clonal selection of the line. These compensatory effects could contribute to the identified transcriptional changes or obscure pathways that are normally regulated by transient pHi changes. To fully understand pH-dependent mechanisms driving normal human epithelial cell biology, we sought to identify the global pH-dependent transcriptome in human epithelial cells subjected to transient and physiological changes in pHi.

Here, we developed an experimental platform to globally identify pH-dependent transcriptional regulation in non-tumorigenic breast epithelial cells (MCF10A). This project improves upon previous work by using small molecule approaches to transiently increase (7.41) and decrease (7.16) pHi of human mammary epithelial cells (7.28) over short (24-hour) timeframes. Our approach avoids the compensatory responses to altered pHi that result from genetic ablation of ion transporters and long-term selection and thus is more likely to recapitulate the transient pHi changes cells experience during normal cell behaviors. We show that just 24 hours of altered pHi triggers a global change in gene expression profiles. After identifying the subset of transcripts that are differentially regulated at high and low pHi, we biochemically validated a collection of pH-dependent hits that clustered in signaling and metabolic pathways.

We show for the first time that the Notch signaling pathway, which is highly conserved in development, is differentially regulated by pHi, with increased Notch1 expression and signaling activity at high pHi compared to control or low pHi. We also show that high pHi transcriptionally upregulates key metabolic enzymes, resulting in altered pyruvate fate and lactate production in normal breast epithelial cells. Importantly, we confirmed these results using two independent pHi manipulation methods and performed confirmatory experiments across various human and canine epithelial cell models. This work advances our understanding of how dynamic pHi regulates transcriptional responses in normal epithelial cells. It also identifies novel pH-dependent pathways and signaling nodes that may underlie normal pH-dependent cell physiology. Finally, this work identifies pH-dependent pathways that may be dysregulated in diseases associated with altered pHi dynamics, such as cancer (increased pHi) (22) and neurodegeneration (lowered pHi) (23).

## Results

### Transient pHi changes in normal mammary epithelial cells do not affect cell viability

We selected MCF10A, an immortalized mammary epithelial cell line, for the initial RNAseq experiments. We note that MCF10A cells do not express estrogen receptors and express high molecular weight keratins (24), which mark MCF10A as having a “basal epithelial” phenotype (25) and not a luminal epithelial phenotype (26). Importantly, MCF10A are non-tumorigenic, near-diploid, genetically stable (27), do not grow in low-adhesion or suspension growth cultures (28), and form single-layer epithelial acini with hollowed out lumens in 3D cell growth (25, 29)— all markers of a robust normal breast epithelial cell model. For these reasons, MCF10A cells are an ideal model of normal human epithelia to characterize the pHi-dependent transcriptome.

We first identified culture conditions that raise and lower pHi in adherent MCF10A cells over 24 hours. For all experiments, we measure pHi at the population level using the cell-permeable pH-sensitive dye 2’,7’-Bis-(2-Carboxyethyl)-5-(and-6)-Carboxyfluorescein, Acetoxymethyl Ester (BCECF-AM) in a plate reader assay as previously described (30, 31). Control MCF10A have a pHi of 7.28±0.01 (mean±SEM), and pHi is significantly reduced (7.16±0.03) after 24 hours of treatment with 25 μM 5-(N-ethyl-N-isopropyl) amiloride (EIPA), a selective inhibitor of the plasma membrane Na^+^-H^+^ exchanger NHE1 (Figure 1B). The pHi of MCF10A can also be significantly increased (7.41±0.02) after 24 hours of treatment with 30 mM ammonium chloride (NH_4_Cl) (Figure 1B). During normal cell behaviors, pHi dynamics of just 0.1-0.2 pHi units are sufficient to alter cell biology(3, 32, 33). These data show we can experimentally lower the pHi of normal epithelial cells into the physiological range observed during hypoxia (34), mitosis (∼7.1-7.2) (3), and in the adult stem cell niche (6.9-7.2) (4). Similarly, we can experimentally increase the pHi of normal cells into the physiological range observed during G2/M transition during the cell cycle (3, 35), migration (1, 36), and metabolic reprogramming (37).

Because cytosolic acidification below 7.0 has been associated with the induction of apoptosis (38), we next assessed cell viability under the pHi-altering conditions using a trypan blue exclusion assay. We avoided redox-dependent viability assays such as CellTiter Blue (39) and mitochondrial activity-based assays (40) because these readouts are sensitive to both metabolic and redox activity changes that may be pHi-dependent (41, 42). There was no significant difference in cell viability between control MCF10A cells and cells with decreased (EIPA) or increased pHi (NH_4_Cl) under these treatment conditions (Figure 1C). Additionally, we did not observe significant abnormalities or changes in MCF10A epithelial colony morphology when pHi was altered for 24 hours (Figure 1D). These results demonstrate that 24-hour treatment with pH manipulation cell culture conditions does not significantly alter viability or colony formation of human epithelial MCF10A cells.

### Transient changes in pHi alter gene expression profiles in normal mammary epithelial cells

Next, we characterized the pH-dependent transcriptional profiles of normal MCF10A cells in response to the short-term changes in pHi (24 h). To do this, we collected RNA from control MCF10A cells and MCF10A cells treated for 24 hours with the pHi manipulation culture conditions described above. We collected six independent biological replicates per treatment condition, prepared RNASeq libraries, and performed NextGen RNASeq to identify pH-dependent transcriptional profiles (Figure 1E). We also performed Principal Component Analysis and observed strong clustering of treatment conditions across biological replicates (Figure S1). All RNASeq transcript data were analyzed compared to control MCF10A, with fold-changes averaged across the six biological replicates (Supplemental Table 1).

From these RNASeq data, we generated volcano plots of transcripts at low pHi (Figure 2A) and high pHi (Figure 2B), where the grey dots indicate transcripts with significant upregulation (log_2_FC > 0.5) or downregulation (log_2_FC < −0.5) compared to control (with FDR < 0.05). We found that low pHi (EIPA) produced 8,660 significantly altered transcripts (4,340 upregulated, 4,320 downregulated) while high pHi (NH_4_Cl) produced 3,529 significantly altered transcripts (1,891 upregulated, 1,638 downregulated) compared to control MCF10A (Figure 2C, Supplemental Table 2). Furthermore, we identified 176 transcripts that exhibited differential expression at low and high pHi (Figure 2C and Figure S2). Of the 176 transcripts with differential expression, 53 transcripts were upregulated at low pHi and downregulated at high pHi, while 123 transcripts were downregulated at low pHi and upregulated at high pHi (Figure 2C and Figure S2). A complete list of differentially regulated genes, fold changes, and significance can be found in Supplemental Table 3. These data show that transient changes in pHi (24 h) can trigger global transcriptional changes in MCF10A cells and that a subset of the transcriptome is differentially regulated by pHi.

**Figure 2.**
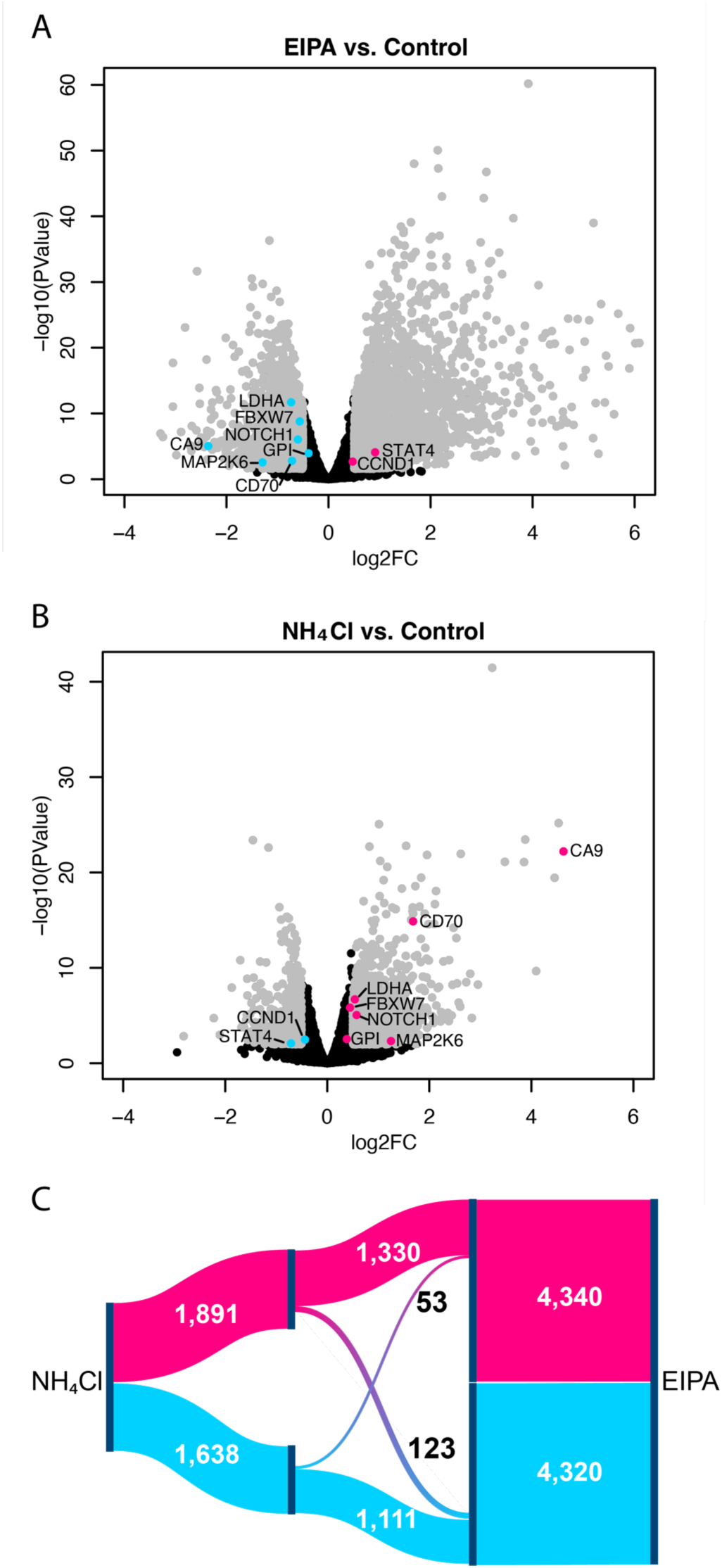
Transient changes in pHi lead to differentially expressed genes in normal epithelial cells. A-B) Volcano plots of RNASeq datasets from RNA collected from normal MCF10A cells treated for 24 hours with (A) EIPA to lower pHi or (B) NH_4_Cl to raise pHi. Shown in grey are transcripts that are upregulated (log_2_FC > 0.5, FDR < 0.05) and downregulated (log_2_FC < −0.5, FDR < 0.5) compared to control. Labeled are a subset of identified transcripts that are differentially regulated at high and low pHi (magenta upregulated, cyan downregulated). C) Sankey diagram showing all significantly altered transcripts from A-B. 176 genes are differentially regulated at high and low pHi, with 53 genes upregulated (magenta) at low pHi and downregulated (cyan) at high pHi, and 123 genes downregulated at low pHi and upregulated at high pHi.

Next, we compared the RNASeq analysis in normal human epithelial cells to the prior microarray dataset obtained from NHE1-null (low pHi) and NHE1-competent (normal pHi) mouse fibroblasts (20). Of the 193 genes reported in the Putney microarray dataset, 73 (37.8%) were also found in the RNAseq dataset of low pHi compared to control (Figure S3A, Supplemental Table 4). Further analysis revealed that of the 73 common genes, 45 (61.6%) had matched transcriptional regulation in both datasets (Figure S3A, Supplemental Table 4), indicating relatively high directionality concordance of pH-sensitive transcripts across the two datasets. However, there were some notable differences between the Putney dataset and our dataset, with 28 genes (38.3%) having mismatched pHi dependence (Figure S3A, Supplemental Table 4). Many of the mismatches fell in metabolic and signaling pathways. For example, the glycolytic enzymes pyruvate kinase muscle isoform (PKM) and hexokinase 2 (HK2) and the cell cycle regulator Wee1 were found to be upregulated at low pHi compared to control pHi in Putney’s mouse fibroblast microarray dataset. However, our RNASeq analysis showed downregulation of these three genes at low pHi compared to control. This lack of complete concordance may indicate that cellular context matters (i.e., human breast epithelial cells vs. mouse fibroblasts) for a subset of pHi-dependent transcripts. Alternatively, the Putney dataset may reflect both pH-dependent effects and compensatory transcriptional effects resulting not from altered pHi but from the genetic ablation, clonal selection, and adaptation required to generate the experimental cell lines used in that work. These compensatory effects would not be present when applying the short-term pHi manipulation approaches we used, which more closely mimic the transient physiological pHi dynamics found in pHi-dependent cell processes (3, 36, 43).

Putney *et al.* identified the most relevant gene ontology terms represented in their microarray dataset from a mouse fibroblast model (20). Interestingly, many of the pathways identified in their study have been shown to have pH-dependent behaviors, such as cell cycle progression (2, 3), carbohydrate metabolism (37, 41, 44), ion transporters (8, 45)(46)(47), and cell adhesion (16, 48). Gene ontology analysis of our pHi-dependent RNASeq dataset from a human epithelial model indicates that signaling (16%) and metabolic pathways (16%) are the two most predominantly altered pathway categories (Figure S3B). These analyses show that transient changes in pHi (24 h) are sufficient to trigger pH-dependent transcriptional regulation in MCF10A and that regulation is clustering in pathways linked to pHi-dependent processes.

We predicted that genes with differential regulation at high pHi compared to low pHi were more likely to indicate a true pH-sensitive node and not an off-target effect of small molecule treatment. We found that differentially-regulated transcripts included pH-homeostasis regulators (Carbonic anhydrase IX, CA9), metabolic proteins (glucose-6-phosphate isomerase, GPI/G6PI; lactate dehydrogenase, LDHA), cell cycle proteins (cyclin D1, CCND1; cyclin K, CCNK), signaling receptors and transcriptional activators (Notch homolog 1, NOTCH1; signal transducer and activator of transcription 4, STAT4), kinases (MAP kinase kinase 6; MAP2K6), and proteasome pathway members (F-box/WD repeat-containing protein 7; FBXW7) (Figure 2A, B). For a complete list of differentially regulated genes, see Figure S2 and Supplemental Table 3. We similarly used MetaCore pathway analysis software (Clarivate Analytics) to identify the cellular pathways with a high incidence of genes differentially regulated by pHi (see methods, Supplemental Table S5). Again, we found that metabolic and signaling pathways were significantly enriched in transcripts differentially regulated by pHi in our RNASeq dataset, with Notch1 being the most significantly enriched signaling pathway and glycolysis being the most significantly enriched metabolic pathway (Figure S3C). Next, we sought to biochemically confirm the pH-dependent expression of Notch1 and glycolytic pathway members.

### NOTCH signaling pathway is upregulated at high pHi and downregulated at low pHi

We used MetaCore pathway analysis to construct a Notch1 signaling pathway based on the RNASeq transcriptional data. Transcripts for NOTCH1 and many downstream pathway members are differentially regulated by pHi (Figure 3A, B). While Notch1 has not been reported to have pH-dependent abundance or function, Notch1 is linked to well-established pH-dependent biology. For example, studies have shown that decreased activity of Notch1 pathway members is associated with stem cell maintenance (49), while upregulation of Notch1 is linked to cell commitment to differentiation in mammary models (50). Stem cell maintenance and differentiation have both been shown to be pH-dependent, with low pHi being required to maintain the stem cell niche and temporal increases in pHi being necessary for differentiation (4). However, despite being associated with these pH-dependent behaviors, Notch1 expression and activity have not been previously shown to be pH-sensitive.

**Figure 3.**
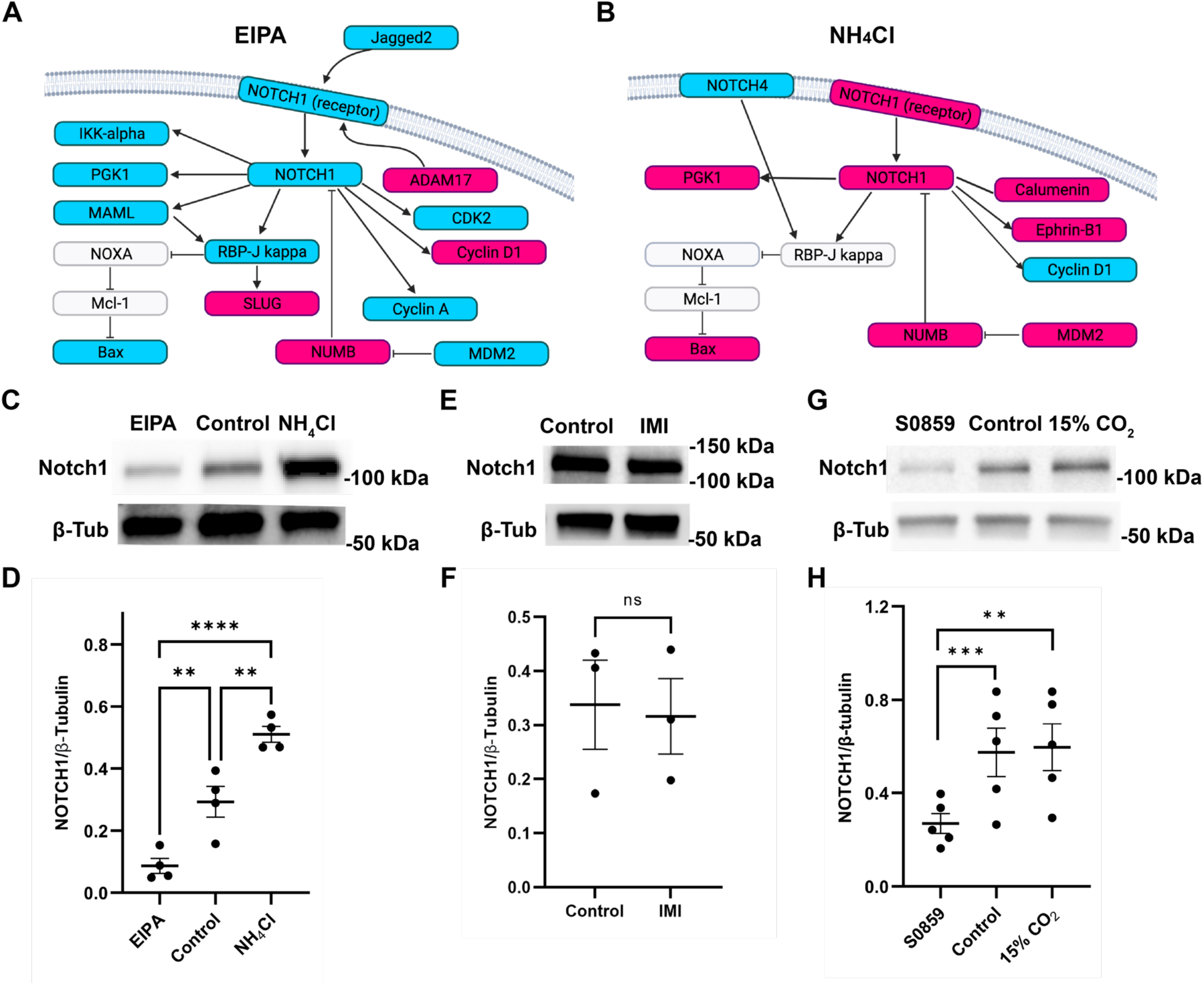
Notch1 expression is increased at high pHi and decreased at low pHi. A-B) Notch1 signaling pathway genes with transcripts that are upregulated (magenta) or downregulated (cyan) at low pHi (EIPA) (A) or high pHi (NH_4_Cl) (B) compared to control MCF10A (Metacore pathway analysis; see methods). C) Representative immunoblot for Notch1 and loading control (β-tubulin; β-Tub) using lysates from control MCF10A cells or MCF10A cells treated for 24 hours with EIPA to lower pHi or NH_4_Cl to raise pHi (see methods for details). D) Quantification of immunoblots prepared as in C. Notch1 expression was normalized to tubulin loading control. Scatter plot (mean±SEM) from 4 biological replicates. E) Representative immunoblot for Notch1 and loading control (β-tubulin; β-Tub) using lysates from control MCF10A cells or MCF10A cells treated for 24 hours with the macropinocytosis inhibitor imipramine (IMI) to ensure the reduction in Notch1 is not due to off-target effects of EIPA inhibition of macropinocytosis. F) Quantification of immunoblots prepared as in E. Notch1 expression was normalized to tubulin loading control. Scatter plot (mean±SEM) from 3 biological replicates. G) Representative immunoblot for Notch1 and loading control (β-tubulin; β-Tub) using lysates from control MCF10A cells or MCF10A cells treated for 24 hours with an inhibitor of the sodium bicarbonate transporter (S0859) to lower pHi or incubated under 15% CO_2_ atmospheric conditions to raise pHi (see methods and Figure S4 for details). H) Quantification of immunoblots prepared as in G. Notch1 expression was normalized to tubulin loading control. Scatter plot (mean±SEM) from 5 biological replicates. For D, significance was determined using ANOVA with Tukey’s multiple comparisons correction. For F and H, significance was determined using a ratio paired t-test. **, p<0.01; ***, p<0.001, ****, p<0.0001.

We found that NOTCH1 is differentially regulated in the RNASeq data (Figure 3A, B; Figure 3A, B). At low pHi, NOTCH1 is downregulated along with several downstream targets of the Notch1 signaling pathway, such as PGK1, CyclinA, CDK2, CBF1, MAML, and IKK alpha (Figure 3A). At high pHi, Notch1 is upregulated, along with several downstream targets of the Notch1 signaling pathway, such as PGK1, Calumenin, and Ephrin-B1 (Figure 3B). Interestingly, we observed that NUMB, a negative regulator of the Notch1 pathway, is upregulated at both high and low pHi, which was initially surprising. However, MDM2, a negative regulator of NUMB, is differentially regulated with high expression at high pHi and low expression at low pHi. The expected outcome of this differential regulation network is an upregulation of the Notch1 signaling pathway at high pHi and downregulation at low pHi. Notch1 is a unique signaling molecule and the pathway lacks an amplification step during signal transduction (51). Thus, Notch1 expression level changes correlate strongly with signal output. Given this pathway result, we predicted that both Notch1 expression and pathway activity would be lower at low pHi and higher at high pHi compared to control MCF10A.

We sought to biochemically validate pH-dependent changes in Notch1 protein abundance and downstream pathway activation. We assayed for protein abundance in MCF10A cell lysates via western blot and found that Notch1 abundance increased at high pHi and decreased at low pHi compared to control MCF10A cells (Figure 3C, D). This biochemical analysis of Notch1 abundance confirms that pH-dependent NOTCH1 transcription leads to pH-dependent Notch1 protein abundance. One reported off-target effect of EIPA is the inhibition of macropinocytosis (52–54). To ensure that the reduction in Notch1 expression at low pHi was the result of lowered pHi and not macropinocytosis inhibition, we treated MCF10A cells with a specific macropinocytosis inhibitor (imipramine) for 24 hours (see methods) and found that imipramine treatment did not affect Notch1 abundance (Figure 3E, F). This indicates that the observed loss of Notch1 was not due to inhibition of macropinocytosis but was instead driven by the lowered pHi driven by EIPA treatment.

To further confirm that Notch1 abundance changes are due to altered pHi in MCF10A and not an off-target effect of the manipulation methods, we developed two orthogonal methods to raise and lower pHi in MCF10A cells. We found that MCF10A pHi is significantly reduced compared to control after 24 hours of treatment with 30 μM S0859, a selective inhibitor of the plasma membrane sodium bicarbonate (Na^+^-HCO_3_^-^) cotransporter(55) (Figure S4). Altering the atmospheric culture conditions to 15% CO_2_ has previously been shown to increase pHi in both normal epithelial models (48) and fibroblasts (16), and we found that the pHi of MCF10A cells was significantly increased (7.63±0.06) after 24 hours at 15% CO_2_ (Figure S4). Using these two orthogonal methods to raise and lower pHi, we observed similar trends in Notch1 abundance, with increased abundance at high pHi compared to low pHi in MCF10A cells (Figure 4G, H). These data show that pH-dependent Notch1 transcriptional changes are sufficient to drive pH-dependent changes in Notch1 protein abundance in MCF10A.

**Figure 4.**
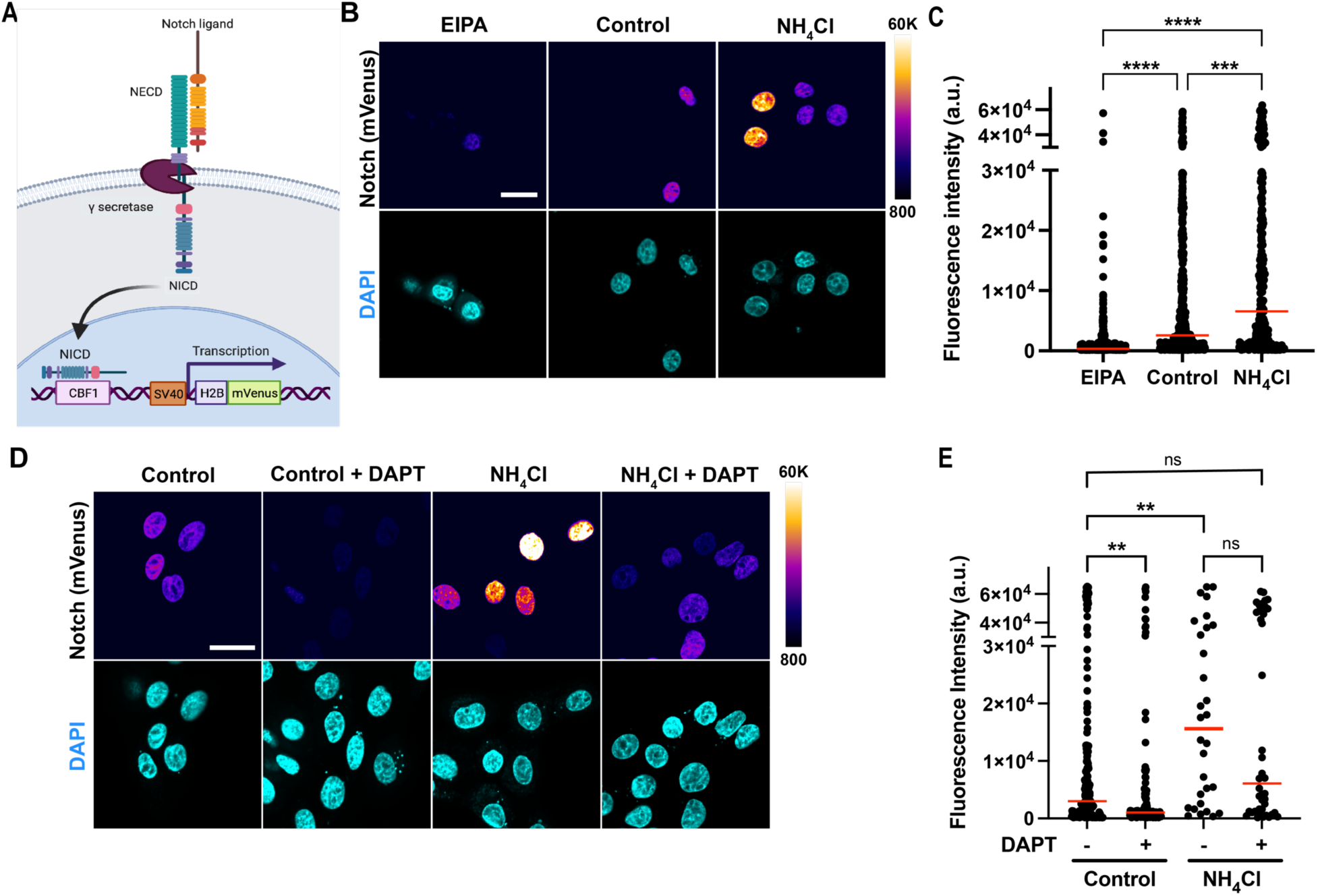
Notch1 transcriptional activity is increased at high pHi and decreased at low pHi. A) Schematic of Notch1 signaling reporter used to quantify Notch1 activity in single-cells. The binding of Notch1 to the CBF promoter drives mVenus fluorescent protein expression. B) Representative confocal images of MCF10A cells expressing the Notch1 activity reporter and treated for 24 hours with EIPA to lower pHi or NH_4_Cl to raise pHi (see methods for details). Notch1 reporter fluorescence is shown in intensiometric display, and nuclei were stained with Hoechst dye (DAPI). Scale bars: 20 μm. C) Quantification of Notch reporter activity from images collected as in B. Reporter fluorescence intensity from individual cell nuclei are shown in a scatter plot (medians shown in red line) from 3 biological replicates. D) Representative confocal images of MCF10A cells expressing the Notch1 activity reporter and treated for 24 hours with NH_4_Cl. Replicate cells were also treated for 24 hours with a gamma-secretase inhibitor (DAPT). Scale bars: 20 μm. E) Quantification of Notch reporter activity from images collected as in D. Reporter fluorescence intensity from individual cell nuclei are shown in a scatter plot (medians shown in red line) from 3 biological replicates. For C and E, significance was determined using Kruskal-Wallis with Dunn’s multiple comparisons correction: **p<0.01, ***p<0.001, ****p<0.0001.

Next, we confirmed pH-dependent Notch1 abundance in two other normal epithelial cell models: human retinal pigment epithelial (RPE) cells and Madin Darby canine kidney (MDCK) cells. We first established experimental pHi manipulation protocols in each cell line. For RPE cells, pHi can be significantly increased using 24 hours of 15% CO_2_ (pHi 7.61±0.02) and significantly decreased using S0859 (pHi 7.33±0.06) (Figure S5A). We found that Notch1 expression in RPE cells was increased ∼2-fold at high pHi compared to low pHi (Figure S5B, C). In MDCK cells, pHi can be increased using 24 hours of incubation at 15% CO_2_ (pHi 7.59±0.16) and decreased using a combination of EIPA and S0859 (pHi 7.14±0.06) (Figure S5D). We found that Notch1 expression in MDCK cells is significantly increased at high pHi and significantly decreased at low pHi compared to control (Figure S5E, F). Our biochemical analyses of protein abundance show that pH-dependent Notch1 expression is conserved across various normal epithelial tissues and species.

We next determined whether pHi-dependent changes in Notch1 abundance are sufficient to alter the transcription of downstream Notch1 target genes. When Notch1 signaling is activated through ligand binding, Notch1 gets cleaved by gamma-secretase at the plasma membrane and the C-terminal domain (NICD) translocates to the nucleus. In the nucleus, Notch1 NICD binds to the CBF/RBP-Jk transcription factor to facilitate binding to the CBF1 promoter and activate transcription of downstream target genes. Thus, to assay for pH-dependent Notch1 pathway activation, we used a Notch1 signaling reporter (56) where nuclear mVenus expression is driven under a CBF1 promoter (Figure 4A). We transfected a large pool of MCF10A cells with the reporter plasmid and then plated cells in triplicate on imaging dishes for 24 hours to recover. The transfected MCF10A cells were then treated with EIPA (low pHi), NH_4_Cl (high pHi), or left untreated (control) for 24 hours before Notch reporter activity was assessed in single cells using confocal microscopy (Figure 4B). We observed that Notch1 signaling activity was increased at high pHi and decreased at low pHi compared to control MCF10A cells (Figure 4C).

To confirm that the increased Notch1 reporter fluorescence at high pHi was due to Notch-dependent transcriptional changes, we repeated these assays with DAPT, a small molecule inhibitor of gamma-secretase that inhibits Notch1 cleavage and translocation to the nucleus(57). We found that DAPT treatment significantly reduced the fluorescence of the Notch1 activity reporter under both high pHi and control conditions (Figure 4D, E). Importantly, DAPT treatment under high pHi conditions reduced the signal of the Notch1 activity reporter to that observed under control conditions (Figure 4E). These controls show that the increases in Notch1 reporter fluorescence observed at high pHi require active gamma-secretase and are thus Notch1-dependent. These data demonstrate that pHi-dependent changes in Notch1 transcript and protein abundance produce pHi-dependent increases in Notch1 pathway activation in normal epithelial cells.

In summary, our RNASeq analyses show that normal mammary epithelial cells exhibit pH-dependent transcription of Notch1 pathway members involved in the cell cycle, metabolism, apoptosis, and proteostasis. Our biochemical and cell biological analyses show that Notch1 protein expression is pH-dependent, leading to pH-dependent Notch1 activity. The Notch signaling pathway is an evolutionarily conserved pathway that has been well characterized for its roles in cellular and tissue development (51), differentiation (50), angiogenesis (58), and proliferation (59). While it has been previously shown that Notch1 signaling is a highly regulated process, including regulation by environmental factors such as temperature(60), this is the first evidence of a role for pHi in regulating Notch1 expression and signaling activity.

### Glycolytic pathway enzymes are upregulated at high pHi and downregulated at low pHi in MCF10A cells

Another pathway that was significantly enriched in genes differentially regulated by pHi was the glycolytic pathway (Figure 5A). Glycolytic pathway enzymes were generally increased at high pHi and decreased at low pHi compared to control MCF10A. Interestingly, our RNASeq data reveal that the expression of both reversible enzymes (such as glucose-6-phosphate isomerase and triosephosphate isomerase) and irreversible enzymes (such as hexokinase and pyruvate kinase) are affected by pHi (Figure 5A).

**Figure 5.**
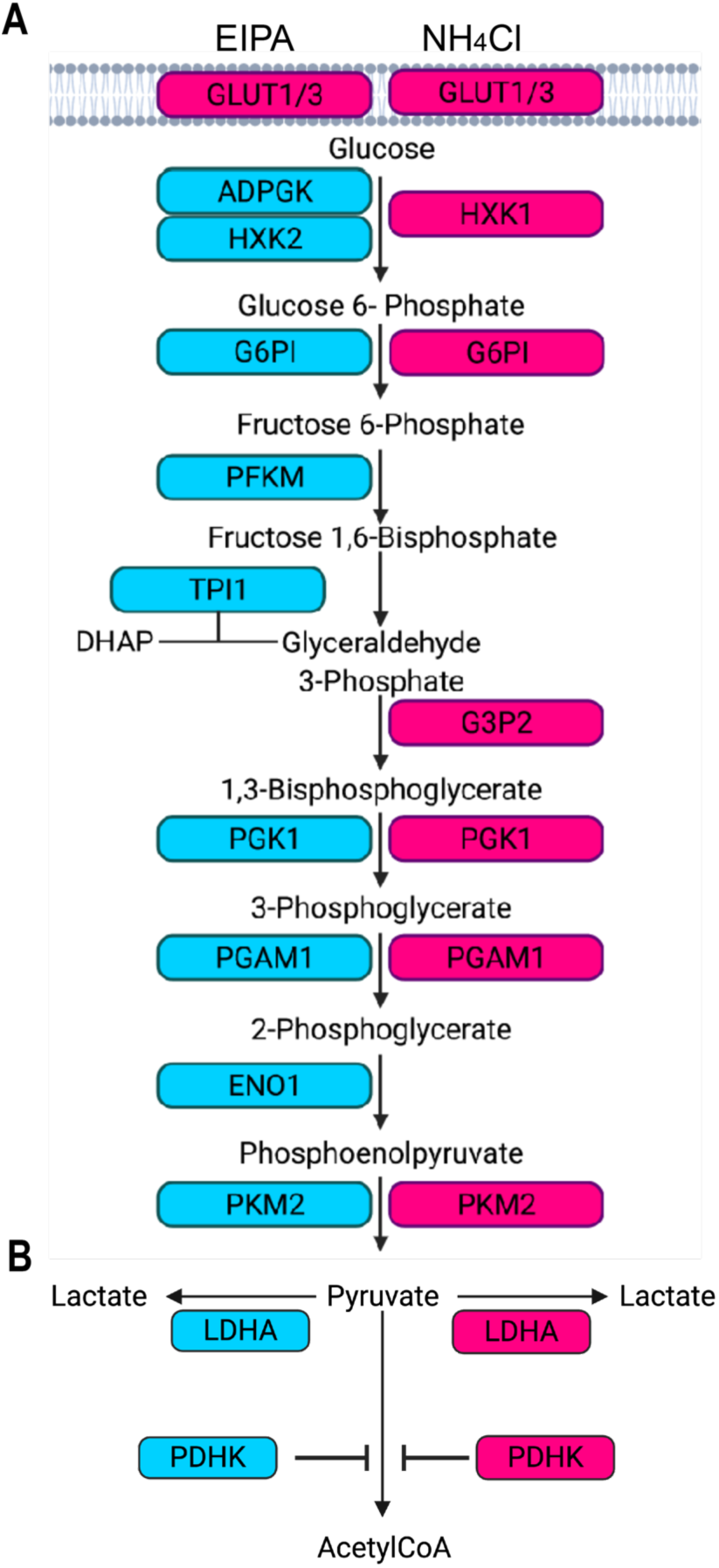
Transcription of glycolytic pathway enzymes and regulators of pyruvate fate are differentially regulated by pHi. A) Glycolytic pathway genes with transcripts that were upregulated (magenta) or downregulated (cyan) at low pHi (EIPA, left) or high pHi (NH_4_Cl, right) compared to the control (MetaCore pathway analysis; see methods). B) Genes that regulate pyruvate fate have transcripts that were upregulated (magenta) or downregulated (cyan) at low pHi (EIPA, left) or high pHi (NH_4_Cl, right).

In addition to these pronounced pH-dependent changes in glycolytic enzyme expression, our RNASeq analysis also revealed pHi-dependent transcription of several enzymes regulating pyruvate fate (Figure 5B). In normal cells, pyruvate is primarily processed through pyruvate dehydrogenase (PDH) to generate Acetyl-CoA (AcCoA) for citric acid cycling and oxidative phosphorylation, while a shift to proliferative (glycolytic) metabolism is marked by conversion of pyruvate to lactate via lactate dehydrogenase A (LDHA) (61). In MCF10A cells, LDHA is differentially regulated by pHi, with upregulation at high pHi and downregulation at low pHi compared to control (Figure 3A). Notably, while LDHA has previously been shown to be downregulated at low pHi (20), matching our observations, our data also reveal that high pHi upregulates LDHA expression compared to control MCF10A cells (Figure 5B). In addition to the transcriptional regulation of LDHA, we also found a novel role for pHi in regulating the expression of pyruvate dehydrogenase kinase (PDK1), which phosphorylates the E1 subunit of PDH to negatively regulate activity and reduce the conversion of pyruvate to AcCoA. PDK1 expression is upregulated at high pHi and downregulated at low pHi compared to control MCF10A (Supplemental Table 3). This suggests that high pHi could increase PDK1-dependent inhibition of PDH and reduce the conversion of pyruvate into AcCoA while simultaneously enhancing LDHA abundance, promoting conversion of pyruvate to lactate. These data indicate that higher pHi inhibits oxidative phosphorylation and could increase glycolytic metabolism. In contrast, low pHi favors decreased PDK1 expression, PDH activation, and high pyruvate conversion into AcCoA for entry into the Krebs cycle during cellular respiration.

### Expression of regulators of pyruvate fate (LDHA and PDK1) are upregulated at high pHi compared to low pHi

Protein abundance does not always correlate with transcript levels, and many glycolytic enzymes are highly allosterically regulated. Thus, we first biochemically confirmed the observed pH-dependent transcriptional changes in LDHA using assays for protein abundance and activity. Using population-level western blot analysis, we observed a trend of higher LDHA protein abundance in MCF10A at high pHi, with significant increases in abundance at high pHi compared to low pHi (Figure S6A, B). We performed single-cell immunofluorescent staining and found that LDHA protein was significantly increased at high pHi and significantly decreased at low pHi compared to control MCF10A (Figure 6A, B). We next confirmed that LDHA expression was pH-dependent in single cells using the previously described orthogonal methods of altering pHi in MCF10A. Again, we found that increased pHi using 15%CO_2_ significantly increased LDHA expression compared to both control MCF10A cells and MCF10A cells where pHi was lowered using S0859 (Figure S7).

**Figure 6.**
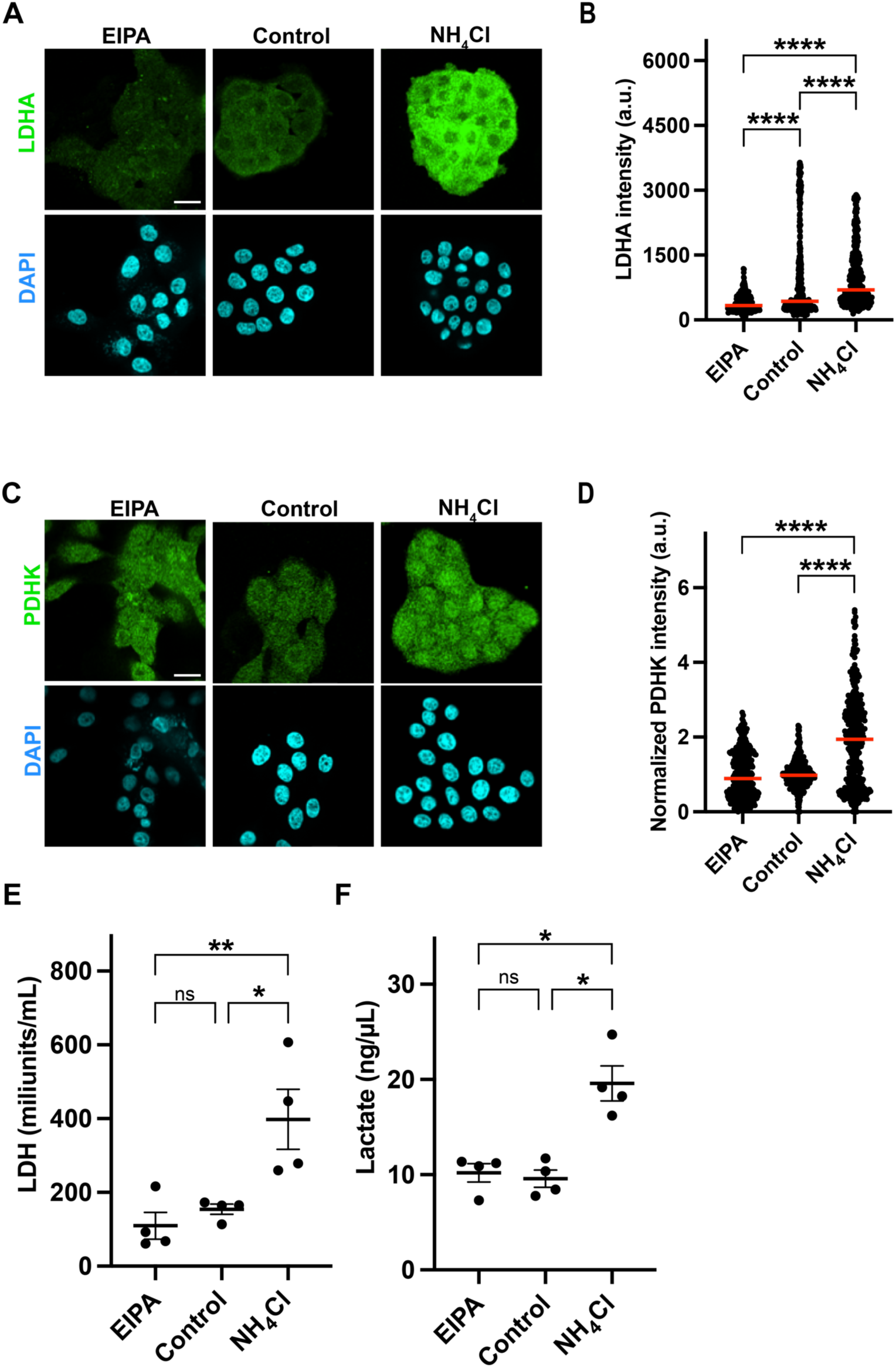
LDHA and pyruvate dehydrogenase kinase expression increase at high pHi and decrease at low pHi in single cells, leading to pH-dependent lactate production. A) Representative confocal images of MCF10A cells treated for 24 hours with EIPA to lower pHi or NH_4_Cl to raise pHi. Cells were then fixed and stained with their corresponding primary and secondary antibodies (see methods). LDHA fluorescence is shown in green, and nuclei stain in cyan (DAPI). Scale bars: 20 μm. B) Quantification of LDHA fluorescent intensity from images collected as in A. Fluorescence intensity from individual cells is shown in a scatter plot (medians shown in red line) from 5 biological replicates. C) Representative confocal images of MCF10A cells treated as in A. Cells were fixed and stained for pyruvate dehydrogenase kinase (PDHK), PDHK fluorescence is shown in green, and nuclei stain in cyan (DAPI). Scale bars: 20 μm. D) Quantification of PDHK fluorescent intensity from images collected as in C. Fluorescence intensity from individual cells are shown in a scatter plot (medians shown in red line) from 3 biological replicates. E) Measurement of LDHA activity in lysates of cells treated as in A. F) Measurement of intracellular lactate in cells treated as in A. Scatter plot (mean±SEM) from 4 biological replicates. For B and D, significance was determined using Kruskal-Wallis with Dunn’s multiple comparisons correction; for E and F, significance was determined using ANOVA with Tukey’s multiple comparisons correction; *p<0.05, **p<0.01, ***p<0.001, ****p<0.0001.

To confirm that pHi-dependent LDHA expression was not unique to MCF10A cells, we also performed single-cell analyses in RPE cells using the previously described methods of raising and lowering pHi (see Fig S5). The data from RPE cells showed similar pH-dependent expression of LDHA protein observed in MCF10A cells, with significantly increased expression at high pHi compared to control and significantly decreased expression at low pHi compared to high pHi (Figure S7 C,D). These results confirm that the observed pH-dependent changes in LDHA transcription produced altered protein abundance in two epithelial models, with increased LDHA abundance at high pHi compared to control and low pHi in both MCF10A and RPE cells.

We next assessed whether transcriptional changes in pyruvate dehydrogenase kinase (PDK1) were sufficient to alter PDK1 protein abundance in MCF10A cells. We found that PDK1 protein abundance was increased at high pHi compared to low pHi and control MCF10A (Figure S6C, D). Again, we performed single-cell immunofluorescent imaging and found that pyruvate dehydrogenase kinase protein (PDHK) was significantly increased in single cells at high pHi when compared to control and low pHi MCF10A cells (Figure 6C, D). These data lead to the hypothesis that increased pHi produces a more glycolytic metabolism by increasing the conversion of pyruvate to lactate through upregulation of LDHA and reducing the conversion of lactate to AcCoA via regulation of PDH activity by PDHK.

### Altered expression of pyruvate fate regulators leads to increased lactate production at high pHi

To test this hypothesis, we conducted two functional assays of LDHA activity in MCF10A. First, we measured the activity of LDH from lysates isolated from control MCF10A cells and MCF10A cells at high and low pHi. We found that the LDH isolated from cells with high pHi was more active than the LDH isolated from cells with control or low pHi (Figure 3E). Next, we measured the concentration of intracellular lactate in control MCF10A cells compared to MCF10A cells with high and low pHi. We found that cellular lactate levels were significantly increased at high pHi compared to both the control and the low pHi conditions; however, no significant difference was found between control and low pHi (Figure 3F). These results demonstrate that changes in pHi can significantly alter the glycolytic pathway at the transcriptional, protein, and functional levels. Additionally, our results support prior work showing that LDHA activity increases with increased pHi (62) and suggests that changes in LDHA transcription and activity could synergize to support cellular metabolism changes.

### Other regulators of pH-dependent biology are differentially regulated by pH

Our data reveal pH-sensitive gene transcription nodes in pathways known to be pHi dependent, such as proliferation and differentiation, but that have no known pH-dependent molecular regulators identified. These results include carbonic anhydrase 9 (CA9), Cyclin D1 (CCND1), and the differentiation regulatory protein CD70 (Figure 2A, B; Supplemental Table 2). CA9 serves to generate extracellular bicarbonate and is upregulated at high pHi and downregulated at low pHi. Upregulation of CA9 is associated with increased proliferation and migration behaviors associated with increased pHi (63, 64). CA9 silencing, on the other hand, suppresses proliferation (65), promotes mitochondrial biogenesis (66), and reduces cell migration (67), all behaviors that are associated with decreased pHi. Our RNASeq data indicate that pHi-dependent regulation of CA9 expression may reinforce these pH-dependent cell processes.

Cyclin D1 (CCND1) is a crucial regulator of cell cycle progression with a distinct role in the transition from G1 to S. Overexpression of CCND1 has been linked to cancer progression by promoting inhibition of the retinoblastoma gene (68), and it has also been shown to promote endocrine resistance to estrogen in breast cancer (69). Additionally, CCND1 has been previously shown to be upregulated at high pHi, leading to proliferation and tumor growth in retinal epithelial cells (70). However, our study shows that CCND1 is downregulated in human mammary epithelial cells at high pHi and upregulated at low pHi (Figure 2A, B; Supplemental Table 2). These findings are supported by prior data showing low pHi upregulates the expression of CCND1 in mouse fibroblasts (20). This result also supports our recent work showing pHi is dynamic during the cell cycle and that low pHi drives cell cycle transitions, including exit from G1 (3). Thus, the transcriptional results presented here will guide our future work characterizing the molecular drivers of pH-sensitive cell cycle progression.

Finally, we found that the cluster of differentiation 70 protein (CD70) is upregulated at high pHi and downregulated at low pHi in our dataset (Figure 2A, B; Supplemental Table 2). High CD70 expression is associated with epithelial-to-mesenchymal transition (EMT) of breast epithelial cells (71), a process associated with increased pHi (21), though no known pH-dependent molecular drivers of EMT are characterized. In addition to the role of CD70 in driving EMT of normal breast epithelia, high CD70 expression is also associated with increased metastatic potential in breast cancer models (72, 73). Our transcriptional results also have the potential to identify putative molecular drivers for pH-dependent EMT, initiation of transformation, and progression of cancer.

### Dataset reveals new putative pH-dependent transcription factors

Our dataset provides the unique opportunity to characterize transcription factors (TFs) with pH-sensitive expression independently from those with predicted pH-sensitive activity. For example, we identified a small number of TFs and co-transcriptional activators with pH-sensitive expression based on the differentially-regulated RNASeq dataset (Figure 7A). This includes TFs with expression that is upregulated at high pHi compared to low pHi (BTG2, E2F7, NOTCH1, TCF19, TP73, ZNF488) as well as TFs with expression that is downregulated at high pHi compared to low pHi (e.g. BEND6, STAT4, and TEF). If pH-dependent transcriptional changes alter TF abundance, downstream signaling of these TFs can be affected, as we show in this work using NOTCH1.

**Figure 7.**
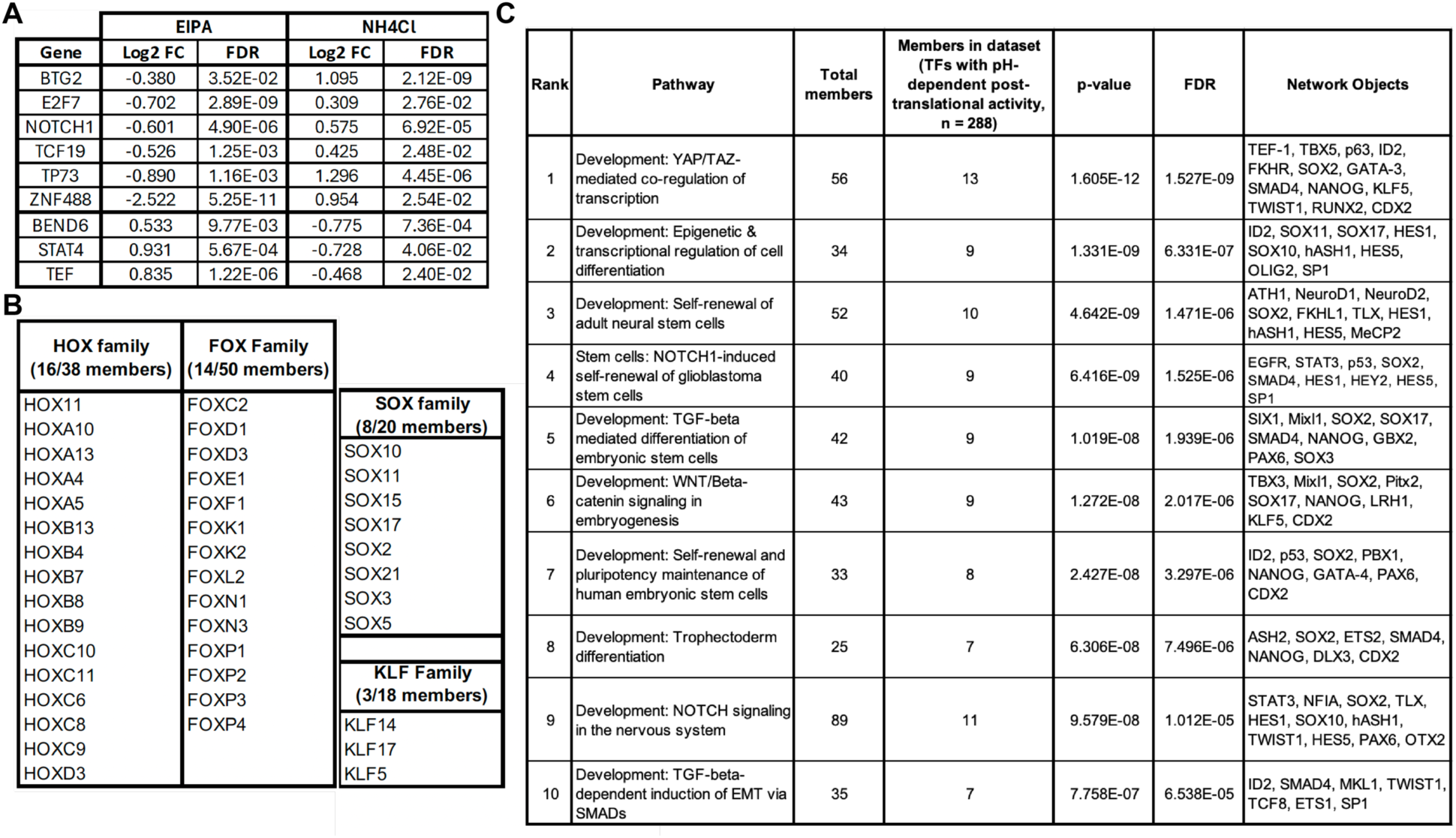
RNAseq dataset can be mined for pH-dependent transcription factor expression and activity. A) Transcription factors (TF) with pH-dependent expression based on the RNASeq dataset. B) TF analysis was performed (see methods) to identify TFs with pH-independent expression but whose downstream targets are pH sensitive. Shown are a subset of these TF families with pH-dependent activity (see Supplemental Table 6 for a complete list of 288 TFs). Families were selected based on recent work showing a role for pHi in directly regulating the DNA binding of these TF families (74). C) TFs in our dataset with pH-dependent activity cluster in developmental and epithelial plasticity pathways. The top 10 pathways are shown here (see Supplemental Table 7 for the top 50 pathways).

However, our dataset also allows us to identify the subset transcription factors with post-translational pH-dependent activity. There are multiple reported modes for pH-dependent post-translational regulation of transcription factor (TF) activity. The first mode is pH-dependent TF stability driving changes in target gene transcription, which we have recently reported for wild-type β-catenin(48) in regulating Wnt signaling in normal epithelial cells(75). The second mode is direct pH-dependent binding of the TF to its target DNA (or co-transcriptional activator), which has been reported previously in mutant p53(31) and in wild-type FOXC2(74). The third mode is pH-dependent regulation of TF subcellular localization, which has been previously reported for wild-type SMAD5(76).

To identify TFs likely to be post-translationally regulated by pHi, we analyzed our RNAseq dataset using the Transcription Factor mode in Metacore (see methods). This approach analyzes the entirety of the RNAseq dataset to determine underlying TF networks based on the end nodes (or targets) of the transcriptionally regulated pathways. Importantly, this algorithm is agnostic to whether the corresponding TF is also present in the original gene list. We filtered these results for TFs not found in our original dataset (i.e. with pHi-independent expression) and identified 288 transcription factors whose downstream targets were significantly and differentially regulated by pHi in our dataset (Supplemental Table 6). Hereafter, we refer to this collection of TFs as TFs with pH-dependent post-translational activity. Importantly, the identified TFs with pH-dependent post-translational activity include TFs previously confirmed to have pH-dependent binding to DNA, including forkhead box transcription factor FOXC2(77) as well as many hits in TF families predicted to be pH-sensitive (based on computational analyses) (78), including HOX family members, SOX family members, and KLF family TFs (Figure 7B). While SMAD5 was not identified in either the dataset of pH-dependent TF expression or pH-dependent TF activity, we did identify SMAD4 in the dataset of TFs with pH-dependent activity (Supplemental Table 6).

We also performed a pathway enrichment analysis on this subset of TFs with pH-dependent post-translational activity and found enrichment of mechanosensitive and developmental pathways (Figure 7C, Supplemental Table 7). This includes pathways linked to pH-dependent differentiation and phenotypic plasticity including: “TGF-B-dependent induction of EMT via SMADs”, which has been shown to be pH-sensitive; Wnt signaling, which we have shown to be pH-sensitive in epithelial cells (75); YAP/TAZ pathways which could contribute to recently characterized pH-dependent mechanosensitive phenotypes(79); and maintenance of STEM cells, which has been shown to be pHi sensitive in intestinal organoids(33). Future work will characterize the molecular mechanisms of these pH-dependent TFs, but our analysis demonstrates the benefit in mining this dataset to rapidly identify putative pH sensitive transcriptional nodes.

## Discussion

Transient changes in pHi enable normal cell behaviors, but in most cases the molecular mechanisms and essential pH-sensitive pathways are not known. Here, we provide evidence that pHi dynamics drive global transcriptional changes in just 24 hours in normal epithelial cells. Our results improve on a previous dataset (20) by using human epithelial cells, increasing transcriptome coverage, and achieving higher specificity of pHi manipulation with reduced cellular adaptation effects. We identified 176 transcripts that are differentially regulated at high pHi compared to low pHi, demonstrating pHi can function as a rheostat in regulating gene transcription in normal epithelial cells. Pathway analysis showed that the top 500 gene ontology (GO) processes affected by changes in pHi were metabolism (16%) and signaling/signal transduction (16%) followed by differentiation, biosynthesis, and cell cycle (with less than 50 GO terms each). This suggests a role for pHi-dependent transcription in directly regulating these pathways. We also validated that pHi drives expression changes in key pathway hubs in the Notch1 signaling pathway (Notch1) and characterized a role for pHi in regulating pyruvate fate inside cells via regulating expression of metabolic pathway enzymes (LDHA and PDK1). Finally, we showed that a subset of TFs involved in development, cell state regulation, and phenotypic plasticity have pHi-dependent post-translational activity.

Our data identify a novel role for increased pHi in driving increased Notch1 signaling in a normal genetic background. Notch1 signaling has been linked to pH-dependent processes like stem cell differentiation (83), epithelial to mesenchymal transition (84), and cell cycle progression (59). This demonstrates that the transient increases in pHi (24 hours) observed during differentiation may drive or reinforce the increased Notch activity required for temporal regulation of differentiation (4, 85, 86). In addition to its role in the normal cell behaviors described above, Notch1 upregulation has been shown to be tumorigenic (87) and associated with epithelial to mesenchymal transition (84). Interestingly, our work establishes a link between these oncogenic roles for Notch1 and the constitutively increased pHi of cancer. Furthermore, these results open the possibility of using pHi to regulate Notch signaling and probe the impact of Notch signaling regulation in distinct biological contexts and processes. We expect that our work will open new lines of research studying pHi regulation of Notch1 as a target to prevent or treat Notch1-related diseases.

Metabolic reprogramming is linked to various normal pH-dependent processes including stem cell differentiation (88), epithelial to mesenchymal transition (EMT) (89), and cell migration (90, 91). However, prior work is conflicting on the role for pHi in directly altering metabolism. Some data indicate that pHi is a driver of metabolic changes (20, 41) while others suggest altered pHi is a response to underlying metabolic changes (92). Additionally, computational analyses have indicated that glycolytic flux may be increased at high pHi and that low pHi may inhibit glycolysis (41). Our data demonstrate that short-term increases in pHi in normal epithelial cells can drive changes in transcript and protein abundances of metabolic enzymes to alter metabolic output of the cells. We show that pHi differentially regulates expression of master regulators of pyruvate fate (such as lactate dehydrogenase and pyruvate dehydrogenase kinase) which serve to integrate pyruvate metabolite flux through various pathways. Thus, our results support pHi as a driving mechanism for metabolic reprogramming and not an outcome of metabolic reprogramming itself.

The pH-dependent transcriptome characterized in this study will have value for epithelial biologists as well as biologists and biochemists working in cancer and neurodegeneration as constitutively increased pHi is a feature of cancer (22, 80, 81), while decreased pHi is linked to neurodegenerative diseases (23). One of the well-established hallmarks of cancer is the metabolic reprograming of cells (93) as they switch from cellular respiration to a glycolytic metabolism. This process is also known as the Warburg effect (94). Our data show that just 24 hours of increased pHi can drive Warburg-like changes in metabolism and support a role for pHi in initiating or reinforcing metabolic changes in tumorigenesis. Our transcriptional results show normal mammary epithelial cells upregulate glycolytic enzymes at high pHi and downregulate glycolytic enzymes at low pHi. Additionally, our biochemical data shows that pHi can differentially regulate pyruvate fate to drive a glycolytic (Warburg-like) metabolism at increased pHi driven by increased expression of LDHA and PDK1 to drive conversion of pyruvate to lactate over AcCoA. Conversely, at the low pHi frequently seen in stem cells with bivalent metabolism (95), pH-dependent loss of LDHA and PDK1 may support preferential conversion of pyruvate to AcCoA.

However, care must be taken not to over-interpret metabolic pathway changes observed in our data as metabolic enzymes are allosterically regulated and metabolic flux is not always correlated to enzyme expression level. Moreover, prior computational approaches have shown that increased pHi is predicted to alter metabolic enzyme activity (41). Persi *et al*. presented a computational prediction for metabolic enzyme activity changes as a function of pH (41). Comparing the Persi results to our expression data, we found that a subset of genes such as CA9, G6PI, and PGK1 are similarly regulated at the transcriptional level in our dataset and proposed and at the enzymatic activity level in their dataset (Supplemental Table 8). Taken together, these data suggest that pHi may play multiple roles in the regulation of enzymatic activity and during the metabolic reprogramming. Future work will collect metabolomic data to support these pHi-dependent transcriptional profiles and integrate the -omics datasets with published computational predicted activity mapping (41).

Our work demonstrates that transient changes in pHi can promote global transcriptional changes in normal epithelial cells. This dataset provides a starting point to explore pHi-associated biomarkers, pH-sensitive pathway nodes, and pH-dependent transcriptional regulation in cell biology. Data presented here show that pHi changes are sufficient drivers of signaling and metabolic changes associated with pH-dependent cell behaviors. Furthermore, our work shows that increased pHi is a sufficient driver of metabolic reprogramming in epithelial cells and supports further study of pHi changes as an early marker of metabolic reprogramming during normal cell processes like differentiation and EMT, as well as the potential for high pHi to drive tumorigenic phenotypes in cancer.

## Acknowledgments

We would like to thank Dr. Joe Sarro, Jacqueline Lopez, and Dr. Michael Pfrender form the Genomics & Bioinformatics Core at the University of Notre Dame for their help in the planning, execution, and analysis of the RNASeq experiments presented in this work. Notch activity reporter plasmid (CBF:H2B-Venus) was a gift from Anna-Katerina Hadjantonakis (Addgene plasmid #44211). This work was supported by an American Cancer Society-Institutional Research Grant (IRG), an NIH New Innovator Award (1DP2CA260416) to K.A.W. as well as startup funds from the University of Notre Dame and the Henry Luce Foundation. The spinning disk confocal microscope used in this work is a part of the Notre Dame Integrated Imaging Facility (NDIIF). Figures 1A, 1E, 3A, 3B, 4A, and 5 were prepared using BioRender software.

## Author Contributions

K.A.W. and R.R.M. conceived the project and planned the experimental strategy. K.A.W. and R.R.M worked in collaboration with the Genomics & Bioinformatics Core at Notre Dame to design the RNASeq strategy. R.R.M., B.J.C., L.M.H, and J.F.K performed the experiments. R.R.M, B.J.C., L.M.H, J.F.K, and K.A.W. analyzed the data. R.R.M., B.J.C., L.M.H, J.F.K, and K.A.W. created the figures. R.R.M., B.J.C., and K.A.W. wrote the first draft manuscript. All authors edited and approved final submission.

## Declaration of interests

The authors declare no competing interests.

## Supplemental Figures and Legends

**Figure S1.**
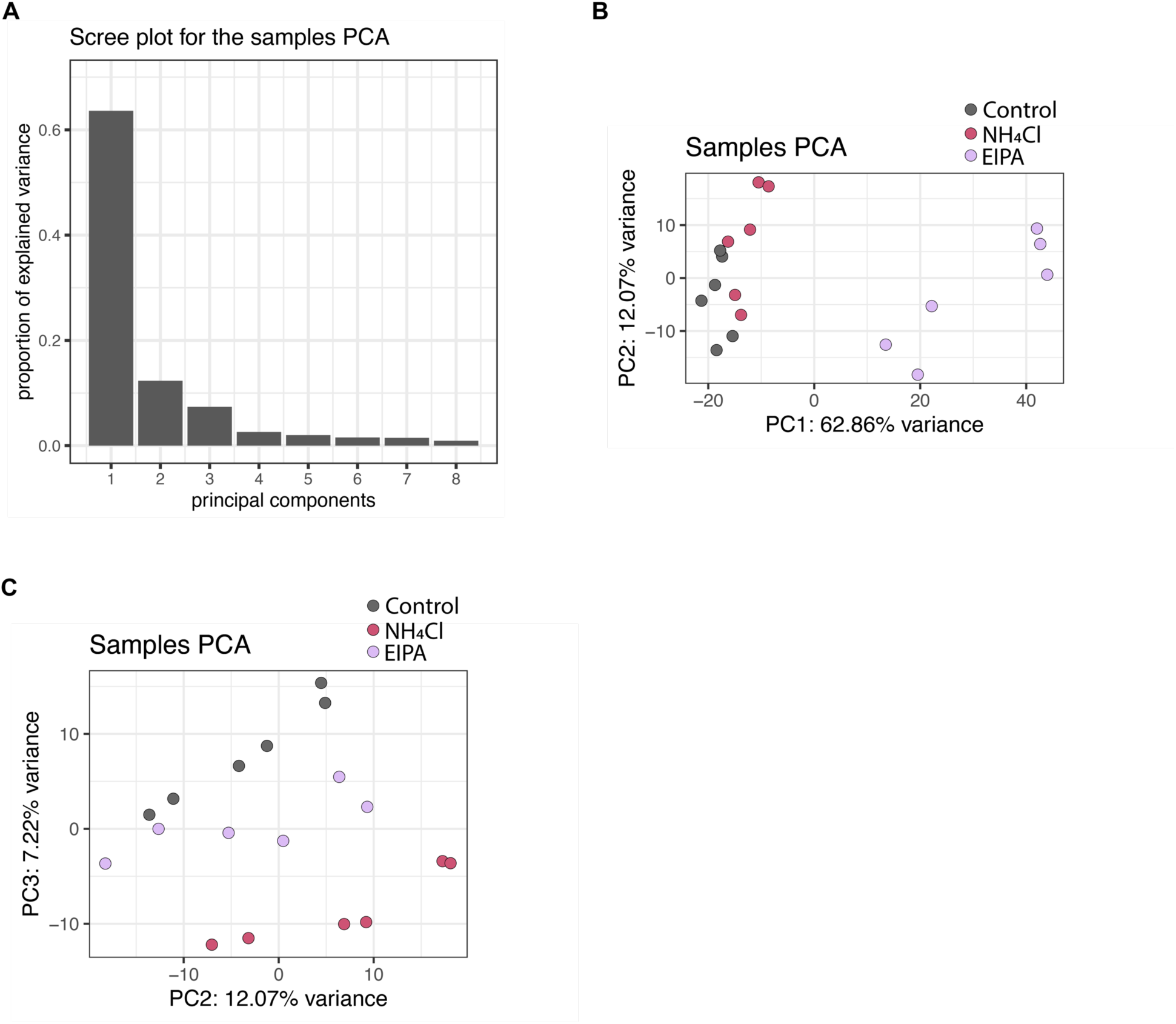
PCA of RNASeq shows that low (EIPA), normal (Control), and high (NH4Cl) pHi MCF10A cells cluster among treatment groups. A) Variance distribution of all principal components calculated in the Principal Component Analysis (PCA) of all 6 RNASeq biological replicates across the 3 different pHi treatment conditions (see methods). Component 1 accounts for just under 63% of the variance, component 2 describes about 12% of the variance, and component 3 describes just over 7% of the total variance in the dataset. B) PCA plot of components 1 and 2 shows a clear separation of low pHi samples from the normal and high pHi conditions. C) PCA plot of components 2 and 3 shows separation between the high pHi samples from the other two conditions.

**Figure S2.**
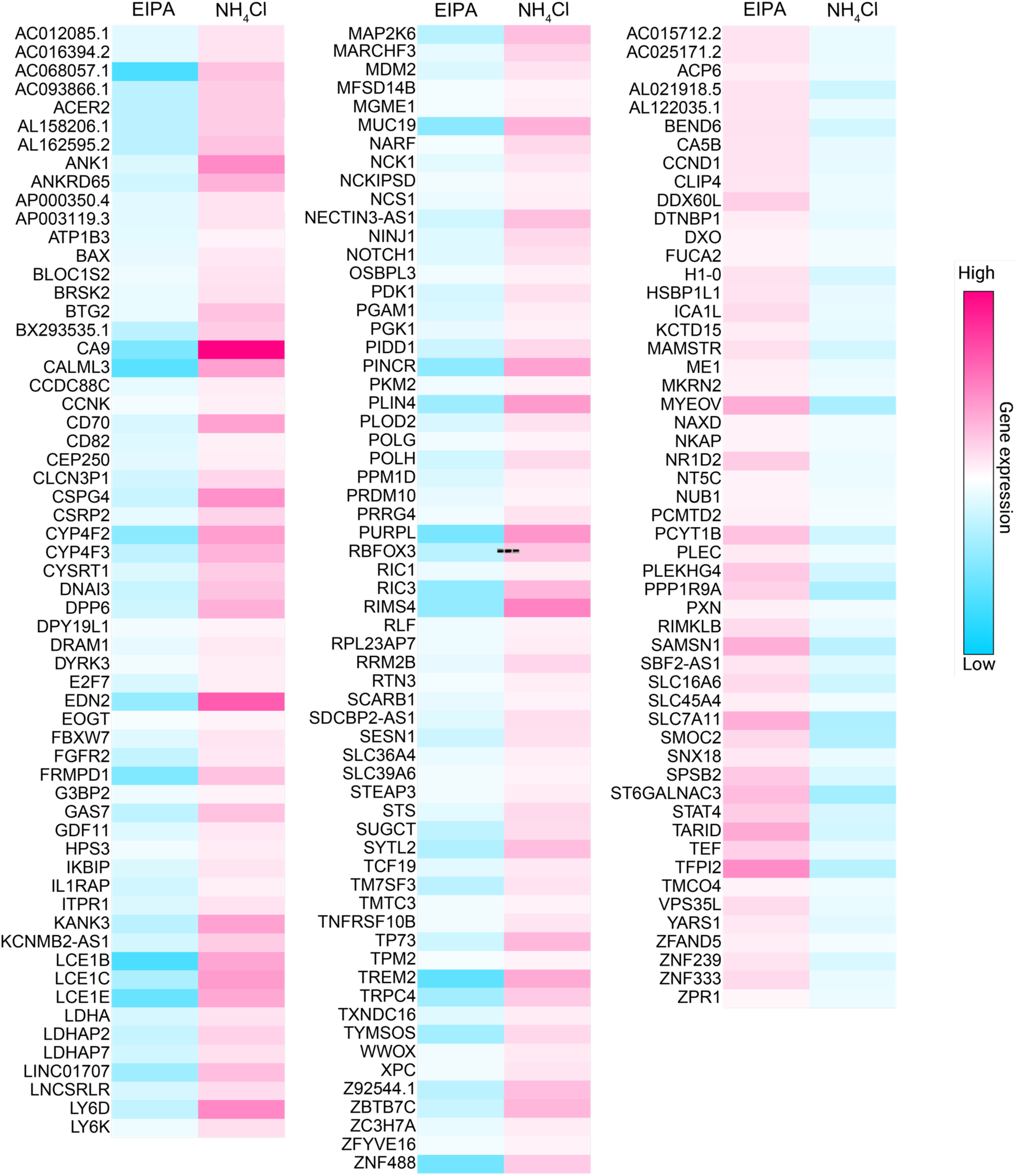
All differentially regulated genes in the RNASeq dataset. Heatmap showing the significantly upregulated (magenta) and downregulated (cyan) genes found in the RNASeq data when compared to the control. Higher opacity corresponds to further deviation from the control sample.

**Figure S3.**
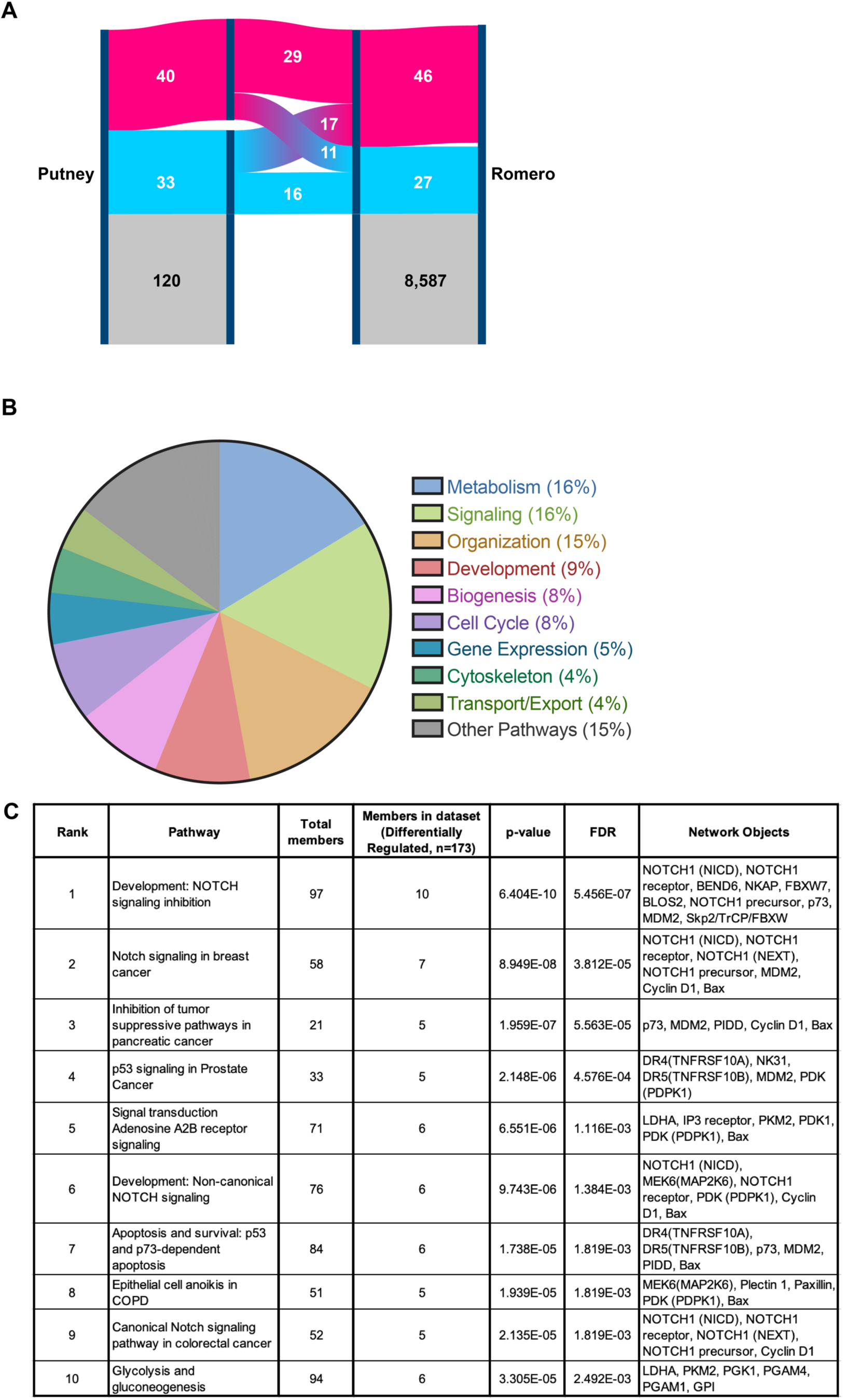
Comparison of gene expression profiles between low pHi (EIPA) and control cells. A) Sankey diagram showing the comparison of the low pHi vs. control mouse fibroblast microarray analysis from Putney et al (Putney and Barber, 2004) and our RNASeq analysis of low pHi vs. control normal human mammary epithelial cells. 45 genes showed similar transcriptional regulation between the datasets and 28 genes showed opposite regulatory effects across the two datasets. Upregulated genes are shown in magenta and downregulated genes are shown in cyan. B) Analysis of the top 500 Gene Ontology (GO) terms found in our RNASeq dataset. (Metacore GO analysis; see methods). All GO terms with 3% or lower representation were grouped as Other Pathways. C) Results of pathway analysis of transcripts differentially regulated by pHi in the RNASeq dataset (see methods; n = 173). Top 10 pathways shown (see Supplemental Table 5 for the top 25 pathways).

**Figure S4.**
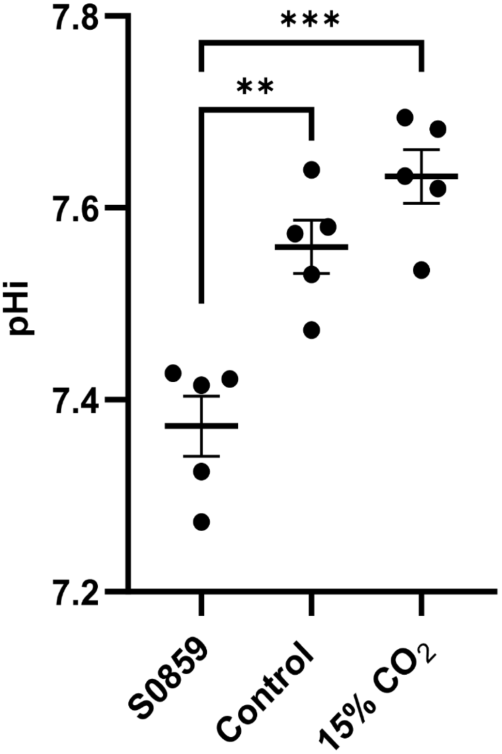
Orthogonal method to manipulate pHi in MCF10A cells. Measured pHi (see methods) in normal human mammary epithelial cells (MCF10A) that were left untreated (control) or treated for 24 hours with complete media containing 30 μM S0859 (S0859) to lower pHi or cultured at 15% CO_2_ (15% CO_2_) to raise pHi. Scatter plot (mean±SEM) from 5 biological replicates. Significance was determined using ANOVA with Tukey’s multiple comparisons correction. **, p<0.01; ***, p<0.0001.

**Figure S5.**
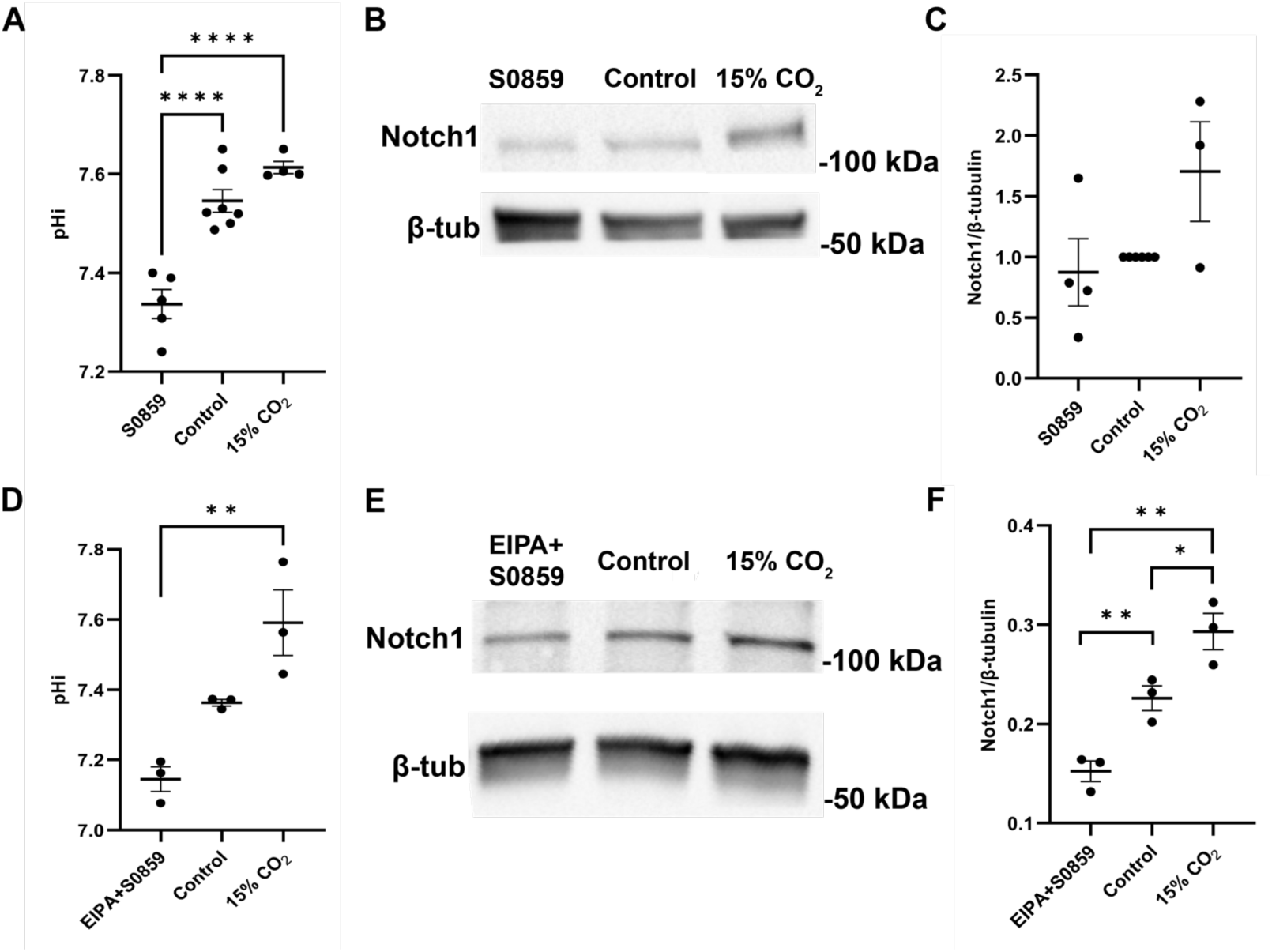
Notch1 expression is pH dependent in other epithelial cell models. A)Retinal pigment epithelial (RPE) cells were treated for 24 hours with 30 μM of S0859 an inhibitor of the sodium bicarbonate transporter to lower pHi and incubated for 24 hours in 15% CO_2_ atmospheric conditions to raise pHi. Scatter plot (mean±SEM) from 5 biological replicates. B) Representative immunoblot for Notch1 and loading control (β-tubulin; β-Tub) using lysates from control RPE cells or RPE cells treated as in A. C) Quantification of immunoblots prepared as in B. Notch1 expression was normalized to tubulin loading control. Scatter plot (mean±SEM) from 4 biological replicates. D) Madin Darby Canine Kidney (MDCK) epithelial cells were treated for 24 hours with 30 μM of S0859 an inhibitor of the sodium bicarbonate transporter and 1 μM of EIPA to lower pHi and incubated for 24 hours in 15% CO_2_ atmospheric conditions to raise pHi. Scatter plot (mean±SEM) from 3 biological replicates. E) Representative immunoblot for Notch1 and loading control (β-tubulin; β-Tub) using lysates from control MDCK cells or MDCK cells treated as in C. F) Quantification of immunoblots prepared as in E. Notch1 expression was normalized to tubulin loading control. Scatter plot (mean±SEM) from 3 biological replicates. Statistical significance was determined using the Kruskal-Wallis test with Dunn’s correction (A), ANOVA with Tukey’s multiple comparisons correction (D) and ratio paired t-test (C,F). *, p<0.05, **, p<0.01, ****, p<0.0001.

**Figure S6.**
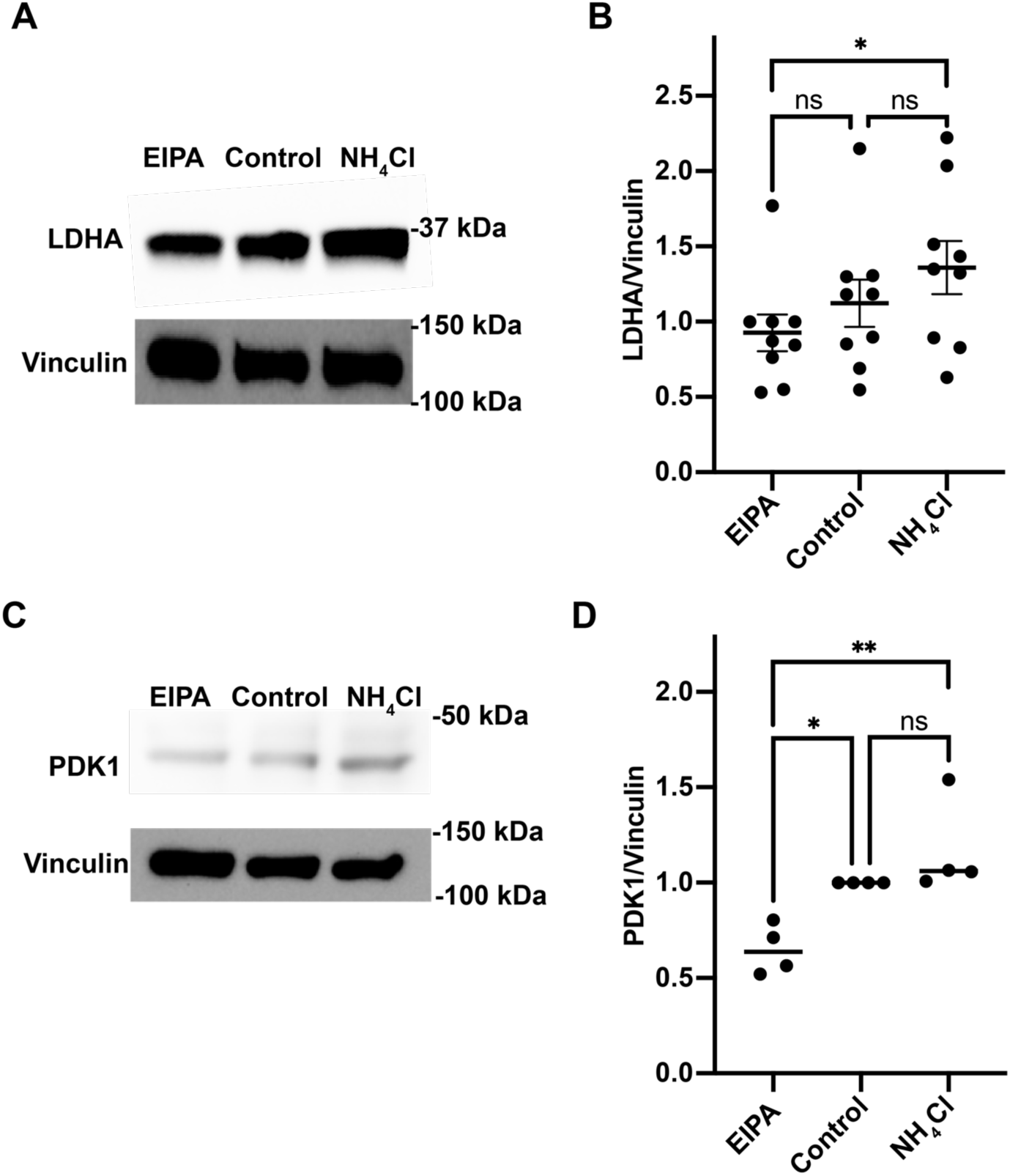
LDHA and PDK1 expression are pH-dependent using population-level western blots to quantify protein abundance. A) Representative immunoblot for lactate dehydrogenase A (LDHA) and loading control (vinculin) using lysates from control MCF10A cells or MCF10A cells treated for 24 hours with EIPA to lower pHi or NH_4_Cl to raise pHi (see methods for details). B) Quantification of immunoblots prepared as in A. LDHA expression was normalized to vinculin loading control. Scatter plot (mean ±SEM) from 9 biological replicates. C) Representative immunoblot for PDK1 and loading control (vinculin) from lysates of MCF10A cells treated as in A. D) Quantification of immunoblots prepared as in C. PDK1 expression was normalized to vinculin loading control. Scatter plot (mean±SEM) from 4 biological replicates. For B and D significance was determined using ANOVA with Tukey’s multiple comparisons correction; *p<0.05, **p<0.01.

**Figure S7.**
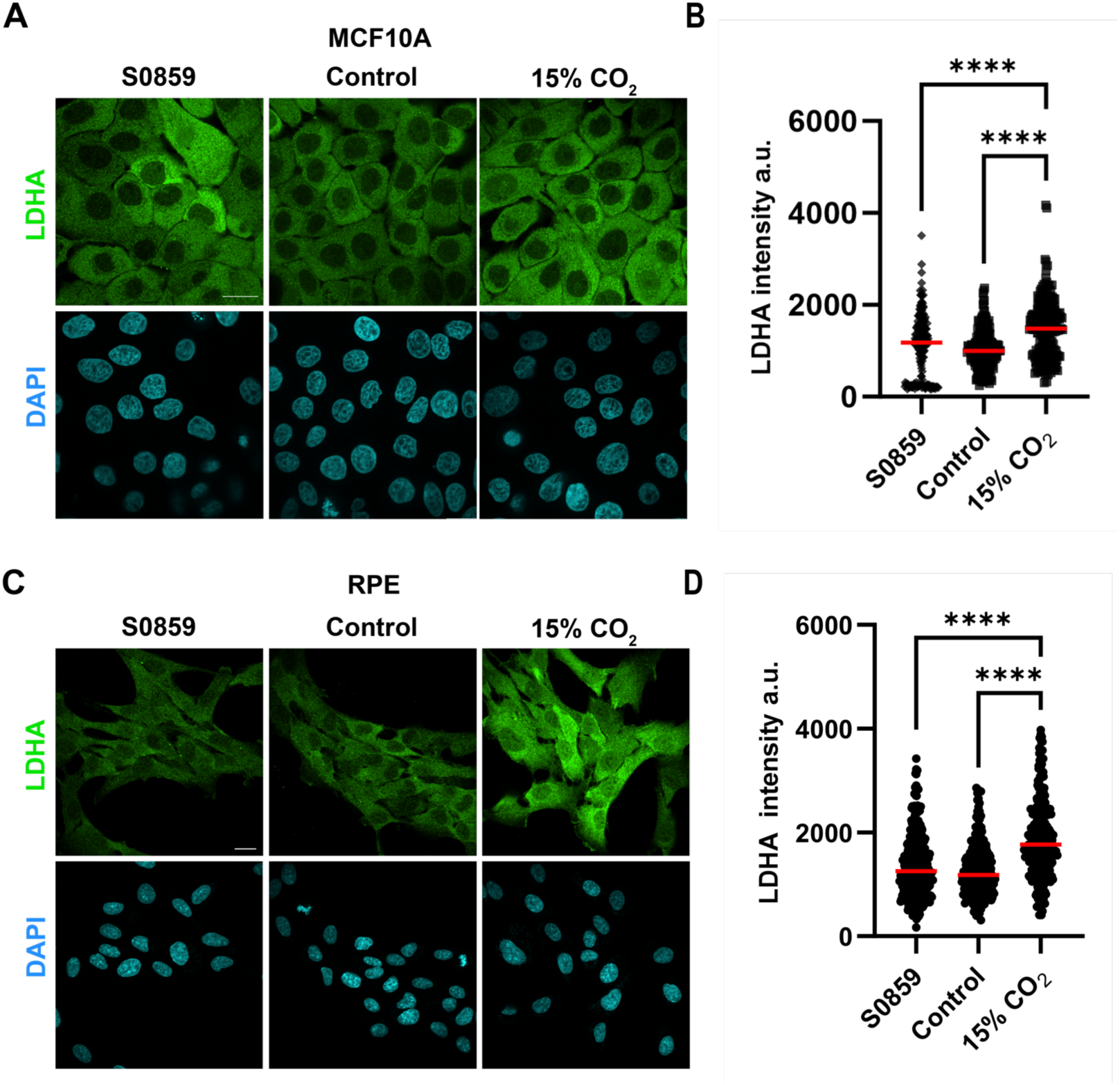
LDHA has pH-dependent expression using orthogonal methods of manipulating pHi and when using another epithelial cell model. A) Representative confocal images of MCF10A cells treated for 24 hours with 30 µM of S0859 an inhibitor of the sodium bicarbonate transporter to lower pHi and incubated for 24 hours in 15% CO_2_ atmospheric conditions to raise pHi. Cells were then fixed and stained. LDHA fluorescence is shown in green and nuclei stain in cyan (DAPI). Scale bars: 20 μm. B) Quantification of LDHA fluorescent intensity from images collected as in A. Fluorescence intensity from individual cells are shown in scatter plot (medians shown in red line) from 5 biological replicates. C) Representative confocal images of RPE cells treated for 24 hours with 30 µM S0859 an inhibitor of the sodium bicarbonate transporter to lower pHi and incubated for 24 hours in 15% CO_2_ atmospheric conditions to raise pHi. Cells were then fixed and stained. LDHA fluorescence is shown in green and nuclei stain in cyan (DAPI). Scale bars: 20 μm. D) Quantification of LDHA fluorescent intensity from images collected as in C. Fluorescence intensity from individual cells are shown in scatter plot (medians shown in red line) from 2 biological replicates. For B and D, significance determined using Kruskal-Wallis with Dunn’s multiple comparisons correction; *p<0.05, **p<0.01, ****p<0.0001.

**Supplemental Table 1. Complete RNASeq Dataset.** All transcripts sequenced and mapped to the human genome in the RNASeq dataset. First tab is all transcripts from low pHi (EIPA) compared to control MCF10A. Second tab is all transcripts from of high pHi (NH4Cl) compared to control. Default sorting is by lowest False Discovery Rate (FDR).

**Supplemental Table 2. All significantly altered transcripts in the RNASeq dataset.** First tab is all transcripts significantly altered at low pHi (EIPA) compared to control MCF10A. Second tab is all transcripts significantly altered at high pHi (NH4Cl) compared to control. Default sorting is alphabetical.

**Supplemental Table 3. All differentially regulated transcripts in the RNASeq dataset.** First tab is all transcripts significantly upregulated at low pHi (EIPA) and downregulated at high pHi (NH_4_Cl) compared to control MCF10A. Second tab is all transcripts significantly downregulated at low pHi (EIPA) and upregulated at high pHi (NH_4_Cl) compared to control MCF10A. Default sorting is alphabetical.

**Supplemental Table 4. Comparison between Putney *et al.* mouse microarray dataset and Romero *et al.* human RNASeq dataset.** First tab is the list of all transcripts found in common between the two datasets and organized as described in Putney *et al.* (Putney and Barber, 2004). Second tab is all transcripts in common listed in alphabetical order. Third tab is only transcripts with matched regulation between the two datasets (concordant data). Fourth tab is only transcripts with opposite regulation between the two datasets (non-concordant data).

**Supplemental Table 5. Pathway Analysis of differentially regulated transcripts in the RNASeq dataset.** List of top 25 pathways enriched in the differentially regulated transcript dataset. Top 10 pathways are shown in Figure S3C.

**Supplemental Table 6. Transcription Factor Analysis of RNASeq dataset.** First tab is the complete transcription factor analysis performed on the RNASeq dataset from MetaCore. All these TFs have downstream targets that are significantly regulated by pHi. Second tab is the subset of TFs that are not in our dataset, but with downstream targets that are significantly regulated by pHi.

**Supplemental Table 7. Pathway analysis of pathways enriched in transcription factors with pH-dependent post-translational activity.** Pathway analysis of the subset of TF genes that aren’t in our dataset, but that have downstream targets that are significantly regulated by pHi. Top 10 are shown in Figure 7C.

**Supplemental Table 8. Comparison between Persi *et al.* computational enzymatic activity prediction and Romero *et al.* human RNASeq dataset.** First tab is transcripts found in common between our RNASeq data set and the predicted behavior of the metabolic enzymes described in Persi *et al.* (Persi *et al.*, 2018). Second tab is only transcripts with matched directionality between the two datasets (concordant data). Third tab is only transcripts with mismatched directionality between the two datasets (non-concordant data).

## Experimental Procedures

### Cell culture

Cells were maintained in humidified incubators at 37°C with 5% CO_2_. Where indicated, atmospheric incubator conditions were altered to 15% CO_2_. MCF10A cells (ATCC: CRL-10317), RPE (ATCC: CRL2302), and MDCK (ATCC: CCL-34) cells were split at sub-confluency and discarded when passaged 30 times after thawing. Cells were monitored for maintenance of epithelial morphology throughout experimental protocols.

MCF10A complete media: DMEM/50% F12 w/ GlutaMax (Invitrogen, 10565-018) supplemented with 5% Horse Serum (Invitrogen, 16050-122), 0.02 ug/mL EGF (Peprotech, AF-100-15), 5 ug/mL Hydrocortisone (Sigma, H-0888), 10 mg/mL Insulin (Sigma, I-1882), 0.1 ug/mL Cholera toxin (Sigma, C-8052), and 1% Penicillin-Streptomycin (Corning, 30-001-Cl).

RPE complete media: RPMI 1640 (Corning, 10-040-CV) supplemented with 10% fetal bovine serum (FBS) (Peak Serum, PS-FB2, lot 06C1182) and GlutaMAX™ (Thermo Fisher, 35050061) for a final glutamine concentration of 446 mg/L.

MDCK complete media: EMEM (with L-glutamine) (Quality Biological, 112-018-101CS) supplemented with 10% FBS (Peak Serum, PS-FB2, lot 06C1182).

pHi manipulation media:

Control: Complete media as described above.

Low pHi: Complete media supplemented with 25 μM EIPA (Invitrogen, E3111) (MCF10A) or 30 µM S0859 (Millipore Sigma, SML0638) (MCF10A, RPE), or a combination of 1 μM EIPA + 30 μM S0859 (MDCK).

High pHi: Complete media supplemented with 30 mM ammonium chloride (NH_4_Cl, Fisher Chemical, A661-500, Lot 185503) (MCF10A) or complete media incubated under 15% CO_2_ conditions (MCF10A, RPE, MDCK).

All chemical concentrations are final concentrations (Cf).

### pHi measurement and calculation

Cells were plated in triplicate for each treatment condition (MCF10: 3×10^4^ cells/well; RPE & MDCK: 4×10^4^ cells/well; 24 well plate; 1 mL total volume). 24 hours after plating, media was removed and pHi manipulation media (see above) was added to the cells. After 24 hours of incubation with pHi manipulation media, pHi was measured as previously described (Choi *et al.*, 2013; White *et al.*, 2017). Briefly, 2’,7’-Bis-(2-Carboxyethyl)-5-(and-6)-Carboxyfluorescein, Acetoxymethyl (BCECF-AM) Ester (Biotium, 51011) was added at 2 μM (Cf) to cells for 30 min at 37°C with 5% CO2 (or 15% CO_2_ for cells treated with 15% CO_2_ to raise pHi). Cells were then washed 3X 5 min with HEPES-based wash buffer (30 mM HEPES, 145 mM NaCl, 5 mM KCl, 10 mM glucose, 1 mM KPO_4_, 1 mM MgSO_4_, and 2 mM CaCl_2_) at 37°C with 5% CO_2_. Wash buffers were supplemented with appropriate pH-altering compounds at the same concentrations as the 24 hr incubations. After washes, fluorescence was read on a plate reader (Molecular Devices Spectramax M3) at the pH-sensitive wavelengths (490ex/535em) and the pH-insensitive wavelengths (440ex/535em) was acquired every 15sec for 5 minutes at Medium PMT sensitivity. After the initial pHi read, wash buffers were removed and replaced with a ∼7.6 pH nigericin buffer standard (25 mM HEPES, 80 mM KCl, 1 mM MgCl_2_) supplemented with 10 μM nigericin (Invitrogen, N1495). Cells were incubated at 37°C for 10 min, and the plate was read again with the same parameters. The high nigericin buffer was then replaced with a ∼6.6 pH nigericin buffer standard, cells were incubated at 37°C for 10 min, and the plate was read again with the same parameters. Mean intensities from pHi measurement and the nigericin standards were exported and pHi was back-calculated using a nigericin linear regression performed on each individual well.

### Trypan blue exclusion assay

Cells were plated at 30,000 cells per well in a 24 well plate. 24 hours after plating, cells were incubated for 24 hours in 30 mM NH_4_Cl, 25 μM EIPA, or fresh media in a 24-well plate. After the 24 h incubation period, the media supernatant was removed and we washed the cells with 2 ml of prewarmed 1xDPBS (Lonza, 17-512F). The PBS was then removed and cells were then incubated at 37°C in 250 μL of trypsin (Corning, 25-053-CI) for about 13-15 minutes until cells started to come off the plate. After incubation in trypsin, 250 μL of prewarmed full culture media was added to each well to inactivate the trypsin. Cells were gently resuspended with care to avoid clumps, then 50 μL of the resuspended cells were added to an Eppendorf tube. Just before counting, 50 μL of trypan blue 0.4% solution (Gibco, 15250-061) was added to the Eppendorf tube and mixed well. 10 μL of the trypan blue and suspended cells mixture was counted on a hemocytometer for healthy (clear) cells and unhealthy/dying (blue) cells. Each biological replicate is the average of three technical replicates (three different wells for each condition). Each biological replicate was assessed from cells plated on a different day. Counts of the multiple conditions were performed in batches to avoid long exposure to trypsin, formation of clumps, or over-incubation in trypan blue.

### RNA extraction and sequencing

From a single parental 10 cm plate of MCF10A cells, when the plate reached a confluency of ∼85%, cells were split 1:6 into three plates. 24 hours later, each one of these three plates was split into two plates that would serve as two technical replicates for each experimental condition (control, 30 mM NH_4_Cl, 25 μM EIPA). After 48 hours following the second passage, media was changed with the appropriate concentration of the pH-altering compounds and incubated at 37°C with 5% CO_2_ for 24 hours. After the 24-hour incubation in pHi-altering treatment, plates were quickly washed 2X with ice cold PBS and then total RNA was extracted following manufacturer’s protocol (RNeasy Plus Minikit (Qiagen, 74134) and RNA concentration and purity was assessed (Thermo Fisher, NanoDrop One).

Sequencing Strategy was developed in collaboration with the Notre Dame Genomics Core: To ensure reproducibility, the RNAseq data was collected across 6 biological replicates and processed thorough QC and library prep in two batches. Eighteen (18) total RNASeq libraries were prepared (three different conditions and six biological replicates per condition) and sequenced across two lanes of an Illumina NextSeq v2.5 (75 cycle) flow cell. Each library was prepared using the NEBNext Poly(A) mRNA Magnetic Isolation Module and NEBNext Ultra II Directional RNA Library Prep kit. QC and quantitation on the library pool were performed using the Qubit dsDNA, Agilent Bioanalyzer DNA 7500 Chip, and Kapa Illumina Library Quantification qPCR assays. Sequencing format was single-read 75bp. Base calling was done by Illumina Real Time Analysis (RTA) v2 software.

Raw sequences were trimmed of adapters with Trimmomatic version 0.39 (Bolger, Lohse and Usadel, 2014) and assessed for quality with FastQC v 0.11.8 (Babraham Bioinformatics). Trimmed sequences were aligned to the human genome, Ensembl build Homo_sapiens.GRCh38, using Homo_sapiens.GRCh38.101 version annotations (Zerbino *et al.*, 2018) and HISAT2 version 2.1.0 (Kim *et al.*, 2019). Corresponding alignments were sorted with SAMtools version 1.9 (Li *et al.*, 2009). Read counts were generated with HTSeq-count version 0.11.2 (Anders, Pyl and Huber, 2015) and were merged with a python script. Subsequent statistics were completed in R (R Core Team, 2014) implementing the EdgeR library (Robinson and Smyth, 2007, 2008; Robinson, McCarthy and Smyth, 2009; McCarthy, Chen and Smyth, 2012). Gene names and GO terms were identified using the Ensembl version of BioMart (Kinsella *et al.*, 2011).

A False Discovery Rate (FDR) < 0.05 cutoff was applied to our RNASeq analysis for pathway analysis and determination of targets for biochemical validation. Log2 fold change (log2FC) < - 0.5 values were considered downregulated, and log2FC > 0.5 values were considered upregulated for each one of our conditions when compared to the control.

### MetaCore Pathway Analysis

After processing the RNASeq data, a table was uploaded to MetaCore containing the mapped gene ID, the log2 Fold Change, and the False Discovery Rate for each comparison (EIPA v. control and NH4Cl v. Control). MetaCore is a manually curated software that builds ontologies mapped to canonical pathways and networks. Once the data was successfully uploaded to MetaCore, the canonical pathway maps analysis was run. This type of analysis results in a list of the pathway maps with the highest z-score for all of the maps in each of our different experimental conditions. The z-score is calculated as follows

z-score = (Actual -Expected) * N * sqrt(N-1) / sqrt(n*R*(N-n)*(N-R)) where Actual is the number of affected genes in a dataset, expected is the mean value for the hypergeometric distribution (n*R/N), N is the total number of nodes on the MetaCore database, n is the total number of nodes in each small network generated from our list, R is the R is the number of the network’s objects corresponding to the genes in our list. And the variance is calculated as follows variance = n*R*(N-n)*(N-R) / (N*N*(N-1)).

This analysis was performed on the rawRNASeq data which was used to generate In addition to performing this analysis on the raw RNASeq data, we also performed the analysis using the subset of differentially regulated genes. This allowed us to find metabolic and signaling pathways that were specifically enriched in differentially regulated genes (see Figure S3C).

### MetaCore Pathway TF Analysis

After processing the RNASeq data, a table was uploaded to MetaCore containing the mapped gene ID, the log2 Fold Change, and the False Discovery Rate for only the statistically significantly different genes from each comparison (EIPA v. control and NH4Cl v. Control). Once the data was successfully uploaded to MetaCore, the Transcription Factor Interactome Analysis was run. This type of analysis shows enrichments of Transcription Factors with the highest z-score in each of our different experimental conditions. The z-score is calculated as described above. Importantly, this analysis is agnostic to whether the TF is in our dataset or not. We then filtered these data for TFs not in our dataset to identify TFs with post-translational pH-sensitive activity.

### Western Blots

From a single parental 10 cm plate of cells, cells were split 1:6 into three 10 cm plates (MCF10A) or counted and plated at a density of 6×10^5^ cells/plate (RPE & MDCK). 24 hours later, the media was changed with the appropriate concentration of the pH-altering compounds and incubated at 37°C with 5% (or 15%) CO_2_ for 24 hours. Where indicated, macropinocytosis was inhibited by treating cells with 10 µM imipramine (Sigma Aldrich, I7393) for 24 hours. After the 24-hour incubation in pHi-altering treatment, plates were quickly washed 2X with ice-cold PBS, and then 500 μL of ice-cold lysis buffer (50 Tris, 150 mM NaCl, 1 mM EDTA, 1 mM DTT, 1 mM Na_3_VO_4_, 1 mM NaF, 1% TritonX-100, and a protease inhibitor cocktail (PierceTM Protease Inhibitor Tablets, A32965; 1 tablet/50 mL of lysis buffer), pH 7.4) was added and plates were incubated on ice for 15 minutes. Cells were then scraped off the plate, transferred into a 1.5 mL Eppendorf tube, and pipetted up and down a couple of times to ensure proper cell membrane lysis. Cell lysates were spun down at max speed in a countertop centrifuge at 4°C for 10 minutes. The clear supernatant was then carefully transferred to a new tube and stored at −80°C until further analysis. Protein concentration was assessed by BCA assay (Thermo Scientific, 34580) using the standard manufacturer protocol. 20 μg of protein lysate was mixed with Laemmli loading buffer (Alfa Aestar, J61337-AC), boiled for five minutes, loaded onto gels (10% Bio-Rad Mini-PROTEAN® TGX Stain-FreeTM Precast Gels, 10026447), and electrophoresed in running buffer (25 mM Tris, 192 mM Glycine, 0.1% SDS) at 100V for 2 hours. Proteins were transferred to a PVDF membrane (Millipore Immobilion-P, IPVH00010) using a wet transfer with freshly made transfer buffer (25 mM Tris, 192 mM Glycine, 20% Methanol) at 60V for 1.5 hours. Before transfer, PVDF membranes were soaked for 5 minutes in 100% methanol and then washed 3X with ddH_2_O. After transferring, membranes were blocked with 5% dry milk (or 5% BSA for phospho-antibodies) in 1x TBST (20 mM Tris, 137 mM NaCl, 0.1% Tween, pH 7.6 with HCl) for one hour and then divided for blotting based on the prestained protein ladder (Bio-Rad, 1610373). Primary antibodies (see antibody table for vendor and dilutions) diluted in 5% BSA were incubated overnight at 4°C followed by 3 x 5-minute washes with 1xTBST and a 1-hour incubation with an HRP-conjugated secondary antibody followed by another 3 x 5-minute wash with 1xTBST. Chemiluminescence was induced by incubating the membranes for 3-5 minutes in the SuperSignalTM West Pico PLUS Chemiluminescent Substrate (Thermo Scientific, 34580). Images were captured through a gel imager (Bio-Rad, ChemiDoc MP).

Western blot analysis was performed using ImageJ densitometry analysis. Each band was enclosed using a rectangular selection of the same size. A densitometry histogram was plotted for the selected experimental and loading control bands. The background was subtracted using the straight-line tool, and the area under the enclosed curve was acquired. The ratio of the protein of interest normalized to loading control was calculated from exported densitometry data.

| Target | Host | Dilution | Company | Cat # | Type | Experiment |
| --- | --- | --- | --- | --- | --- | --- |
| NOTCH1 | Rabbit | 1:1000,<br>1:400 | Cell<br>Signaling | 3608S | Primary | WB, IF |
| PDK1 | Rabbit | 1:1000 | Cell<br>Signaling | 3062 | Primary | WB, IF |
| LDHA | Rabbit | 1:1000 | Cell<br>Signaling | 3582S | Primary | WB |
| p-LDHA | Rabbit | 1:1000 | Cell<br>Signaling | 8176S | Primary | WB |
| B-Tubulin | Rabbit | 1:1000 | Abcam | ab6046 | Primary | Loading<br>control |
| Vinculin | Rabbit | 1:1000 | Cell<br>Signaling | 13901S | Primary | Loading<br>control |
| Anti-mouse<br>Alexa 647 | Goat | 1:1000 | Invitrogen | A21236 | Secondary | IF |
| Anti-rabbit<br>Alexa 488 | Goat | 1:1000 | Invitrogen | A11008 | Secondary | IF |
| Anti-mouse<br>HRP | Goat | 1:1000 | Southern<br>Biotech | 1030-05 | Secondary | WB |
| Anti-rabbit<br>HRP | Goat | 1:1000 | Southern<br>Biotech | 1050-05 | Secondary | WB |

### Microscopy

All imaging experiments were performed using a Nikon Ti2 Eclipse confocal microscope equipped with a spinning disk (CSU-X1, Yokogawa) using solid-state lasers (395 nm, 488 nm, 561 nm) with appropriate filter sets (DAPI: ET455/50M, mVENUS/AF488: ET525/36M, AF561: ET561/60M) on 40x objective (CFI Plan Fluor 40X, NA =1.30, MRH01401, NIKON), using a CMOS camera (ORCA-Flash4.0). Cells were imaged in 35 mm imaging dishes (Matsunami, Dd35-14-1.5-U) within a stage-top environmental chamber (Tokai) at 37°C and 5% CO_2_. All microscope control and image analyses used Nikon NIS Elements AR software.

### Notch activity assay

Notch activity reporter (CBF:H2B-Venus) was a gift from Anna-Katerina Hadjantonakis (Addgene plasmid #44211) (Nowotschin *et al.*, 2013). MCF10A cells were grown in a 10 cm dish until they reached 70-80% confluency and then were transfected with CBF:H2B-Venus using manufacturer’s protocol (FuGene HD, Promega, E2311). Briefly, 450 μL of room temperature serum-free media were added to a 0.6 mL Eppendorf tube, and 10 μg of DNA was then added to the Eppendorf tube and mixed by inverting the tube 4-5 times. Then, 30 μL of the transfection reagent (Fugene HD) was added to the tube (3:1 Fugene HD to DNA ratio, μL:μg), mixed again by inverting the tube 4-5 times and then let incubate at room temperature for 15 minutes. After the 15-minute incubation, the contents of the Eppendorf tube were combined with 4.5 mL of pre-warmed full media, and then the mixture was added to the 10 cm plate of cells and incubated at 37°C. Cells were incubated with transfection media for 6 hours and then trypsinized (0.25% Trypsin EDTA, Corning, 25-053-Cl), plated in triplicate (25,000 cells/dish; 35 mm imaging dishes: Matsunami, Dd35-14-1.5-U), and then returned to the incubator overnight. 24 hours after transfection, dishes were treated with pHi manipulation media (see above) with or without the gamma-secretase inhibitor (2*S*)-*N*-[(3,5-Difluorophenyl)acetyl]-L-alanyl-2-phenyl]glycine 1,1-dimethylethyl ester (DAPT; Tocris, 2634) at a final concentration of 10 µM. 48 hours post-transfection (24 hours after pHi manipulation), dishes were imaged by confocal microscopy. A minimum of 10 fields of view were taken using a GFP laser (488 nm excitation, 400 ms, 25% power), a DAPI laser (405 nm excitation, 300 ms, 15% power), and a DIC image (300 ms) using a 40x oil immersion objective. Images were processed and analyzed using Nikon Elements AR software. Images were background subtracted using an ROI placed on an area without cells within each field of view. ROIs were drawn on each cell using the mVenus fluorescence signal, and average ROI intensity values were obtained for each cell’s mVenus signal. To account for artifactual fluorescence, a separate 10 cm plate of cells was mock transfected (same protocol as above but without DNA) and imaging was conducted as described above for all of our conditions. Two times the average fluorescence in these mock-transfected cells was used as a cutoff for inclusion in the analysis.

### LDH assay

The lactate dehydrogenase colorimetric assay (Abs450) kit (Sigma, MAK066) measures LDH activity in a biological sample based on the reduction of NAD to NADH. The amount of NADH generated is used to back-calculate the milliunits/mL of LDH in the biological sample using a linear regression of known standards. For each condition, 1×10^6^ cells were lysed in 500 μL of the assay buffer provided in the kit, and 2 μL of the lysed cells were used for analysis. The manufacturer’s standard protocol was followed for the rest of the analysis. Assays were run in technical duplicates and averaged for each data point.

### Lactate assay

The lactate assay kit (Sigma, MAK066) is a colorimetric assay that measures lactate concentration (0.2–10 nmoles of lactate) in a biological sample and is determined by an enzymatic assay leading to a readout that is proportional to the lactate present in the sample. The lactate concentration in the biological sample can be back-calculated from the read measurement (Abs570)/fluorometric (λex535/ λem587) using a linear regression of known standards. For each condition, 1×10^6^ cells were lysed in 500 μL of the assay buffer provided in the kit. After lysing the cells, samples were spun at 10,000 x g for 10 minutes in a 10 kDa size exclusion tube (EMD Millipore, UFC801024) to remove proteins, with the flow-through retained for further analysis. 2 μL of the extracted cell lysate was used for analysis. The manufacturer’s standard protocol was used for the rest of the analysis. Assays were run in technical duplicates and averaged for each data point.

### Immunofluorescence Assays

Cells were trypsinized and plated (80,000 cells/dish, MCF10A; 50,000 cells/dish, RPE) in separate 3.5 cm imaging dishes for each condition (Matsunami, Dd35-14-1.5-U) and returned to the incubator for 24 hours. Cells were then treated with pHi-manipulation media, and dishes were returned to the incubator overnight. 24 hours after treatment, cells were rinsed with PBS and fixed in 3.7% formaldehyde (Alfa Aestar, 33314) in DPBS at room temperature for 10 minutes. Cells were washed 3×2 min with DPBS, then incubated in antibody buffer (0.1% Triton-X, 1.0% BSA in DPBS) for 10 minutes at room temperature. Cells were washed 3 x 2 min with DPBS, then incubated in blocking buffer (1.0% BSA in DPBS) for 1 hour at RT with rocking followed by 3 x 2 min wash in DPBS. Cells were then incubated overnight at 4°C in primary antibodies (see antibody table for vendor/cat and dilutions) in antibody buffer. After incubation with primary antibody, cells were washed 3 x 2 min in DPBS, then incubated with secondary antibodies (see antibody table for vendor/cat and dilutions), Phalloidin conjugate, and 1:20,000 dilution of Hoescht dye in antibody buffer for one hour at room temperature in the dark with gentle rocking. After a final 3 x 2 min wash in PBS, cells were imaged using a Nikon Ti2 Eclipse confocal microscope equipped with a spinning disk microscope (CSU-X1, Yokogawa). A minimum of 5 fields of view per dish were acquired using the following settings: (AF488: 10% laser power, 500 ms; AF647: 10% laser power, 500 ms; DAPI: 10% laser power, 150 ms), and a DIC image (50 ms). The acquired images were then analyzed using the Nikon NIS Elements AR software: A rectangular ROI on the glass coverslip was used to subtract the background from the current frame in each field of view. Bezier ROIs were used to select each cell as a region of interest. ROIs were saved, and data with GFP intensity values for each ROI was exported into Excel. These data were imported into GraphPad Prism for analysis. After collecting all biological replicates, the ROUT (Q=1%) method was used to identify and remove outliers and Shapiro-Wilk normality tests run in PRISM 9. For non-normally distributed data, a Kruskal-Wallis test with Dunn’s multiple comparisons test was used to identify significant differences between the EIPA, control, and NH_4_Cl. The mean rank of each column was compared with the mean rank of every other column.

### Statistical analysis summary

For all data consisting of 10 data points or more, a Shapiro-Wilk normality test was used to determine the normality of the data distribution. For normally distributed data or data samples smaller than 10 data points, statistical significance was determined using a one-way ANOVA with Tukey multiple-comparisons correction. Normalized immunoblot data was analyzed using a ratio paired t-test between low and high pHi conditions and a one-sample t-test for comparing low and high pHi to control (hypothetical mean = 1.0). For non-normally distributed data, statistical significance was determined using Kruskal−Wallis test with Dunn’s multiple-comparisons correction.

## Data Availability

All necessary data are available in the submitted manuscript or supplemental materials. Any additional data can be requested from the corresponding author.

## FIGURE PREPARATION DETAILS/SOFTWARE

### PCA/METACORE/SANKEY

Figures 1A, 1E, 3A, 3B, 4A, and 5 were prepared using BioRender.com

Figures 1B, 1C, 3D, 3F, 3H, 4C, 4E, 6B, 6D-F, S4, S5A, S5C-D, S5F, S6B, S6D, S7B, and S7D were prepared using GraphPad PRISM

Figures 1D, 4B, 4D, 6A, 6C, S7A and S7C were prepared using the Nikon NIS Elements AR software

Figures 2A, 2B, S1A, S1B, S1C and S2 were created using R/RStudio or PCAExplorer package for R and modified in Adobe Illustrator

Figures 2C and S3A were created using sankeymatic.com and modified in Adobe Illustrator. Figure S3B was created using Microsoft Excel and modified in Adobe Illustrator Metacore (Clarivate) was used to analyze RNASeq datasets. From these analyses, we generated simplified pathways including only genes contained in our dataset for Figure 3A, 3B, and 5.

## Notes

### Competing Interest Statement

The authors have declared no competing interest.

### Summary of Updates

We have added new biochemical and cell biological characterization of pH-dependent transcriptional regulation including using orthogonal methods to manipulate pHi and validating our data in additional normal epithelial cell lines. We have also added new bioinformatic characterization of pH-dependent transcription factor regulation. Text is significantly revised for clarity and detail.

## References

1. Denker, S. P., and Barber, D. L. (2002) Cell migration requires both ion translocation and cytoskeletal anchoring by the Na-H exchanger NHE1. J Cell Biol. 159, 1087–96

2. Putney, L. K., and Barber, D. L. (2003) Na-H exchange-dependent increase in intracellular pH times G2/M entry and transition. J Biol Chem. 278, 44645–9

3. Spear, J. S., and White, K. A. (2023) Single-cell intracellular pH dynamics regulate the cell cycle by timing the G1 exit and G2 transition. Journal of Cell Science. 136, jcs260458

4. Ulmschneider, B., Grillo-Hill, B. K., Benitez, M., Azimova, D. R., Barber, D. L., and Nystul, T. G. (2016) Increased intracellular pH is necessary for adult epithelial and embryonic stem cell differentiation. J Cell Biol. 215, 345–355

5. Halestrap, A. P., and Wilson, M. C. (2012) The monocarboxylate transporter family—Role and regulation. IUBMB Life. 64, 109–119

6. Alka, K., and Casey, J. R. (2014) Bicarbonate transport in health and disease: Bicarbonate Transporters in Health and Disease. IUBMB Life. 66, 596–615

7. Collins, M. P., and Forgac, M. (2020) Regulation and function of V-ATPases in physiology and disease. Biochimica et Biophysica Acta (BBA) - Biomembranes. 1862, 183341

8. Putney, L. K., Denker, S. P., and Barber, D. L. (2002) The Changing Face of the Na ^+^ /H ^+^ Exchanger, NHE1: Structure, Regulation, and Cellular Actions. Annu. Rev. Pharmacol. Toxicol. 42, 527–552

9. Al-Samir, S., Papadopoulos, S., Scheibe, R. J., Meißner, J. D., Cartron, J.-P., Sly, W. S., Alper, S. L., Gros, G., and Endeward, V. (2013) Activity and distribution of intracellular carbonic anhydrase II and their effects on the transport activity of anion exchanger AE1/SLC4A1: Role of CAII in the function of AE1. The Journal of Physiology. 591, 4963–4982

10. Vince, J. W., and Reithmeier, R. A. (2000) Identification of the carbonic anhydrase II binding site in the Cl(-)/HCO(3)(-) anion exchanger AE1. Biochemistry. 39, 5527–33

11. Schonichen, A., Webb, B. A., Jacobson, M. P., and Barber, D. L. (2013) Considering protonation as a posttranslational modification regulating protein structure and function. Annu Rev Biophys. 42, 289–314

12. Stoeckelhuber, M., Noegel, A. A., Eckerskorn, C., Köhler, J., Rieger, D., and Schleicher, M. (1996) Structure/function studies on the pH-dependent actin-binding protein hisactophilin in *Dictyostelium* mutants. Journal of Cell Science. 109, 1825–1835

13. Srivastava, J., Barreiro, G., Groscurth, S., Gingras, A. R., Goult, B. T., Critchley, D. R., Kelly, M. J., Jacobson, M. P., and Barber, D. L. (2008) Structural model and functional significance of pH-dependent talin-actin binding for focal adhesion remodeling. Proc Natl Acad Sci U S A. 105, 14436–41

14. Frantz, C., Barreiro, G., Dominguez, L., Chen, X., Eddy, R., Condeelis, J., Kelly, M. J., Jacobson, M. P., and Barber, D. L. (2008) Cofilin is a pH sensor for actin free barbed end formation: role of phosphoinositide binding. J Cell Biol. 183, 865–79

15. Frantz, C., Karydis, A., Nalbant, P., Hahn, K. M., and Barber, D. L. (2007) Positive feedback between Cdc42 activity and H+ eflux by the Na-H exchanger NHE1 for polarity of migrating cells. Journal of Cell Biology. 179, 403–410

16. Choi, C. H., Webb, B. A., Chimenti, M. S., Jacobson, M. P., and Barber, D. L. (2013) pH sensing by FAK-His58 regulates focal adhesion remodeling. J Cell Biol. 202, 849–59

17. Suzuki, A., Maeda, T., Baba, Y., Shimamura, K., and Kato, Y. (2014) Acidic extracellular pH promotes epithelial mesenchymal transition in Lewis lung carcinoma model. Cancer Cell International. 10.1186/s12935-014-0129-1

18. Cardone, R. A., Alfarouk, K. O., Elliott, R. L., Alqahtani, S. S., Ahmed, S. B. M., Aljarbou, A. N., Greco, M. R., Cannone, S., and Reshkin, S. J. (2019) The Role of Sodium Hydrogen Exchanger 1 in Dysregulation of Proton Dynamics and Reprogramming of Cancer Metabolism as a Sequela. Int J Mol Sci. 10.3390/ijms20153694

19. Alfarouk, K. O., Verduzco, D., Rauch, C., Muddathir, A. K., Adil, H. H., Elhassan, G. O., Ibrahim, M. E., David Polo Orozco, J., Cardone, R. A., Reshkin, S. J., and Harguindey, S. (2014) Glycolysis, tumor metabolism, cancer growth and dissemination. A new pH-based etiopathogenic perspective and therapeutic approach to an old cancer question. Oncoscience. 1, 777–802

20. Putney, L. K., and Barber, D. L. (2004) Expression profile of genes regulated by activity of the Na-H exchanger NHE1. BMC Genomics. 5, 46

21. Amith, S. R., Wilkinson, J. M., and Fliegel, L. (2016) Na+/H+ exchanger NHE1 regulation modulates metastatic potential and epithelial-mesenchymal transition of triple-negative breast cancer cells. Oncotarget. 7, 21091–113

22. White, K. A., Grillo-Hill, B. K., and Barber, D. L. (2017) Cancer cell behaviors mediated by dysregulated pH dynamics at a glance. J Cell Sci. 130, 663–669

23. Harguindey, S., Reshkin, S. J., Orive, G., Arranz, J. L., and Anitua, E. (2007) Growth and trophic factors, pH and the Na+/H+ exchanger in Alzheimer’s disease, other neurodegenerative diseases and cancer: new therapeutic possibilities and potential dangers. Curr Alzheimer Res. 4, 53–65

24. Perou, C. M., Jeffrey, S. S., Van De Rijn, M., Rees, C. A., Eisen, M. B., Ross, D. T., Pergamenschikov, A., Williams, C. F., Zhu, S. X., Lee, J. C. F., Lashkari, D., Shalon, D., Brown, P. O., and Botstein, D. (1999) Distinctive gene expression patterns in human mammary epithelial cells and breast cancers. Proc. Natl. Acad. Sci. U.S.A. 96, 9212–9217

25. Debnath, J., Muthuswamy, S. K., and Brugge, J. S. (2003) Morphogenesis and oncogenesis of MCF-10A mammary epithelial acini grown in three-dimensional basement membrane cultures. Methods. 30, 256–268

26. Prat, A., and Perou, C. M. (2009) Mammary development meets cancer genomics. Nat Med. 15, 842–844

27. Soule, H. D., Maloney, T. M., Wolman, S. R., Peterson, W. D., Brenz, R., McGrath, C. M., Russo, J., Pauley, R. J., Jones, R. F., and Brooks, S. C. (1990) Isolation and characterization of a spontaneously immortalized human breast epithelial cell line, MCF-10. Cancer Res. 50, 6075–6086

28. Puleo, J., and Polyak, K. (2021) The MCF10 Model of Breast Tumor Progression. Cancer Research. 81, 4183–4185

29. Debnath, J., Mills, K. R., Collins, N. L., Reginato, M. J., Muthuswamy, S. K., and Brugge, J. S. (2002) The Role of Apoptosis in Creating and Maintaining Luminal Space within Normal and Oncogene-Expressing Mammary Acini. Cell. 111, 29–40

30. Sesanto, R., Kuehn, J. F., Barber, D. L., and White, K. A. (2021) Low pH Facilitates Heterodimerization of Mutant Isocitrate Dehydrogenase IDH1-R132H and Promotes Production of 2-Hydroxyglutarate. Biochemistry. 60, 1983–1994

31. White, K. A., Garrido Ruiz, D., Szpiech, Z. A., Strauli, N. B., Hernandez, R. D., Jacobson, M. P., and Barber, D. L. (2017) Cancer-associated arginine-to-histidine mutations confer a gain in pH sensing to mutant proteins. Science Signaling. 10, eaam9931

32. Parks, S. K., and Pouyssegur, J. (2015) The Na(+)/HCO3(-) Co-Transporter SLC4A4 Plays a Role in Growth and Migration of Colon and Breast Cancer Cells. J Cell Physiol. 230, 1954–63

33. Liu, Y., Reyes, E., Castillo-Azofeifa, D., Klein, O. D., Nystul, T., and Barber, D. L. (2023) Intracellular pH dynamics regulates intestinal stem cell lineage specification. Nat Commun. 14, 3745

34. Hulikova, A., Harris, A. L., Vaughan-Jones, R. D., and Swietach, P. (2013) Regulation of intracellular pH in cancer cell lines under normoxia and hypoxia. Journal Cellular Physiology. 228, 743–752

35. Andersen, A. P., Flinck, M., Oernbo, E. K., Pedersen, N. B., Viuff, B. M., and Pedersen, S. F. (2016) Roles of acid-extruding ion transporters in regulation of breast cancer cell growth in a 3-dimensional microenvironment. Mol Cancer. 15, 45

36. Martin, C., Pedersen, S. F., Schwab, A., and Stock, C. (2011) Intracellular pH gradients in migrating cells. American Journal of Physiology-Cell Physiology. 300, C490–C495

37. Alfarouk, K. O., Ahmed, S. B. M., Elliott, R. L., Benoit, A., Alqahtani, S. S., Ibrahim, M. E., Bashir, A. H. H., Alhoufie, S. T. S., Elhassan, G. O., Wales, C. C., Schwartz, L. H., Ali, H. S., Ahmed, A., Forde, P. F., Devesa, J., Cardone, R. A., Fais, S., Harguindey, S., and Reshkin, S. J. (2020) The Pentose Phosphate Pathway Dynamics in Cancer and Its Dependency on Intracellular pH. Metabolites. 10.3390/metabo10070285

38. Lagadic-Gossmann, D., Huc, L., and Lecureur, V. (2004) Alterations of intracellular pH homeostasis in apoptosis: origins and roles. Cell Death Differ. 11, 953–61

39. Fields, R. D., and Lancaster, M. V. (1993) Dual-attribute continuous monitoring of cell proliferation/cytotoxicity. Am Biotechnol Lab. 11, 48–50

40. Mosmann, T. (1983) Rapid colorimetric assay for cellular growth and survival: Application to proliferation and cytotoxicity assays. Journal of Immunological Methods. 65, 55–63

41. Persi, E., Duran-Frigola, M., Damaghi, M., Roush, W. R., Aloy, P., Cleveland, J. L., Gillies, R. J., and Ruppin, E. (2018) Systems analysis of intracellular pH vulnerabilities for cancer therapy. Nat Commun. 9, 2997

42. Zhao, H., Zhang, Y., Pan, M., Song, Y., Bai, L., Miao, Y., Huang, Y., Zhu, X., and Song, C.-P. (2019) Dynamic imaging of cellular pH and redox homeostasis with a genetically encoded dual-functional biosensor, pHaROS, in yeast. Journal of Biological Chemistry. 294, 15768– 15780

43. Donahue, C. E. T., Siroky, M. D., and White, K. A. (2021) An optogenetic tool to raise intracellular pH in single cells and drive localized membrane dynamics, Biochemistry, 10.1101/2021.03.09.434608

44. Dietl, K., Renner, K., Dettmer, K., Timischl, B., Eberhart, K., Dorn, C., Hellerbrand, C., Kastenberger, M., Kunz-Schughart, L. A., Oefner, P. J., Andreesen, R., Gottfried, E., and Kreutz, M. P. (2010) Lactic acid and acidification inhibit TNF secretion and glycolysis of human monocytes. J Immunol. 184, 1200–9

45. Webb, B. A., White, K. A., Grillo-Hill, B. K., Schönichen, A., Choi, C., and Barber, D. L. (2016) A Histidine Cluster in the Cytoplasmic Domain of the Na-H Exchanger NHE1 Confers pH-sensitive Phospholipid Binding and Regulates Transporter Activity. J. Biol. Chem. 10.1074/jbc.M116.736215

46. Gorbatenko, A., Olesen, C. W., Boedtkjer, E., and Pedersen, S. F. (2014) Regulation and roles of bicarbonate transporters in cancer. Front Physiol. 5, 130

47. Counillon, L., Bouret, Y., Marchiq, I., and Pouyssegur, J. (2016) Na(+)/H(+) antiporter (NHE1) and lactate/H(+) symporters (MCTs) in pH homeostasis and cancer metabolism. Biochim Biophys Acta. 1863, 2465–80

48. White, K. A., Grillo-Hill, B. K., Esquivel, M., Peralta, J., Bui, V. N., Chire, I., and Barber, D. L. (2018) β-Catenin is a pH sensor with decreased stability at higher intracellular pH. The Journal of Cell Biology. 217, 3965–3976

49. Micchelli, C. A., and Perrimon, N. (2006) Evidence that stem cells reside in the adult Drosophila midgut epithelium. Nature. 439, 475–479

50. Bouras, T., Pal, B., Vaillant, F., Harburg, G., Asselin-Labat, M.-L., Oakes, S. R., Lindeman, G. J., and Visvader, J. E. (2008) Notch Signaling Regulates Mammary Stem Cell Function and Luminal Cell-Fate Commitment. Cell Stem Cell. 3, 429–441

51. Andersson, E. R., Sandberg, R., and Lendahl, U. (2011) Notch signaling: simplicity in design, versatility in function. Development. 138, 3593–3612

52. Koivusalo, M., Welch, C., Hayashi, H., Scott, C. C., Kim, M., Alexander, T., Touret, N., Hahn, K. M., and Grinstein, S. (2010) Amiloride inhibits macropinocytosis by lowering submembranous pH and preventing Rac1 and Cdc42 signaling. J. Cell Biol. 188, 547–563

53. Ivanov, A. I. (2008) Pharmacological Inhibition of Endocytic Pathways: Is It Specific Enough to Be Useful? in Exocytosis and Endocytosis (Ivanov, A. I. ed), pp. 15–33, Methods in Molecular Biology, Humana Press, Totowa, NJ, 440, 15–33

54. West, M. A. (1989) Distinct endocytotic pathways in epidermal growth factor-stimulated human carcinoma A431 cells [published erratum appears in J Cell Biol 1990 Mar;110(3):859]. The Journal of Cell Biology. 109, 2731–2739

55. Ch’En, F. F., Villafuerte, F. C., Swietach, P., Cobden, P. M., and Vaughan-Jones, R. D. (2008) S0859, an *N* -cyanosulphonamide inhibitor of sodium-bicarbonate cotransport in the heart. British J Pharmacology. 153, 972–982

56. Nowotschin, S., Xenopoulos, P., Schrode, N., and Hadjantonakis, A.-K. (2013) A bright single-cell resolution live imaging reporter of Notch signaling in the mouse. BMC Dev Biol. 13, 15

57. Dong, Z., Huo, J., Liang, A., Chen, J., Chen, G., and Liu, D. (2021) Gamma-Secretase Inhibitor (DAPT), a potential therapeutic target drug, caused neurotoxicity in planarian regeneration by inhibiting Notch signaling pathway. Science of The Total Environment. 781, 146735

58. Siekmann, A. F., and Lawson, N. D. (2007) Notch Signalling and the Regulation of Angiogenesis. Cell Adhesion & Migration. 1, 104–105

59. Wang, X.-D., Leow, C. C., Zha, J., Tang, Z., Modrusan, Z., Radtke, F., Aguet, M., De Sauvage, F. J., and Gao, W.-Q. (2006) Notch signaling is required for normal prostatic epithelial cell proliferation and differentiation. Developmental Biology. 290, 66–80

60. Nomura, T., Nagao, K., Shirai, R., Gotoh, H., Umeda, M., and Ono, K. (2022) Temperature sensitivity of Notch signaling underlies species-specific developmental plasticity and robustness in amniote brains. Nat Commun. 13, 96

61. Vander Heiden, M. G., Cantley, L. C., and Thompson, C. B. (2009) Understanding the Warburg Effect: The Metabolic Requirements of Cell Proliferation. Science. 324, 1029–1033

62. Read, J. A., Winter, V. J., Eszes, C. M., Sessions, R. B., and Brady, R. L. (2001) Structural basis for altered activity of M- and H-isozyme forms of human lactate dehydrogenase. Proteins. 43, 175–85

63. Chiche, J., Ilc, K., Laferrière, J., Trottier, E., Dayan, F., Mazure, N. M., Brahimi-Horn, M. C., and Pouysségur, J. (2009) Hypoxia-Inducible Carbonic Anhydrase IX and XII Promote Tumor Cell Growth by Counteracting Acidosis through the Regulation of the Intracellular pH. Cancer Research. 69, 358–368

64. Benej, M., Svastova, E., Banova, R., Kopacek, J., Gibadulinova, A., Kery, M., Arena, S., Scaloni, A., Vitale, M., Zambrano, N., Papandreou, I., Denko, N. C., and Pastorekova, S. (2020) CA IX Stabilizes Intracellular pH to Maintain Metabolic Reprogramming and Proliferation in Hypoxia. Front. Oncol. 10, 1462

65. Cianchi, F., Vinci, M. C., Supuran, C. T., Peruzzi, B., De Giuli, P., Fasolis, G., Perigli, G., Pastorekova, S., Papucci, L., Pini, A., Masini, E., and Puccetti, L. (2010) Selective Inhibition of Carbonic Anhydrase IX Decreases Cell Proliferation and Induces Ceramide-Mediated Apoptosis in Human Cancer Cells. J Pharmacol Exp Ther. 334, 710–719

66. Xu, J., Zhu, S., Xu, L., Liu, X., Ding, W., Wang, Q., Chen, Y., and Deng, H. (2020) CA9 Silencing Promotes Mitochondrial Biogenesis, Increases Putrescine Toxicity and Decreases Cell Motility to Suppress ccRCC Progression. IJMS. 21, 5939

67. Csaderova, L., Debreova, M., Radvak, P., Stano, M., Vrestiakova, M., Kopacek, J., Pastorekova, S., and Svastova, E. (2013) The effect of carbonic anhydrase IX on focal contacts during cell spreading and migration. Front. Physiol. 10.3389/fphys.2013.00271

68. Direct binding of cyclin D to the retinoblastoma gene product (pRb) and pRb phosphorylation by the cyclin D-dependent kinase CDK4 (1993) Genes & Development. 7, 331–342

69. Hui, R., Finney, G. L., Carroll, J. S., Lee, C. S. L., Musgrove, E. A., and Sutherland, R. L. (2002) Constitutive overexpression of cyclin D1 but not cyclin E confers acute resistance to antiestrogens in T-47D breast cancer cells. Cancer Res. 62, 6916–6923

70. Koch, L. M., Birkeland, E. S., Battaglioni, S., Helle, X., Meerang, M., Hiltbrunner, S., Ibáñez, A. J., Peter, M., Curioni-Fontecedro, A., Opitz, I., and Dechant, R. (2020) Cytosolic pH regulates proliferation and tumour growth by promoting expression of cyclin D1. Nat Metab. 2, 1212–1222

71. Sigurdsson, V., Hilmarsdottir, B., Sigmundsdottir, H., Fridriksdottir, A. J. R., Ringnér, M., Villadsen, R., Borg, A., Agnarsson, B. A., Petersen, O. W., Magnusson, M. K., and Gudjonsson, T. (2011) Endothelial Induced EMT in Breast Epithelial Cells with Stem Cell Properties. PLoS ONE. 6, e23833

72. Liu, L., Yin, B., Yi, Z., Liu, X., Hu, Z., Gao, W., Yu, H., and Li, Q. (2018) Breast cancer stem cells characterized by CD70 expression preferentially metastasize to the lungs. Breast Cancer. 25, 706–716

73. Petrau, C., Cornic, M., Bertrand, P., Maingonnat, C., Marchand, V., Picquenot, J.-M., Jardin, F., and Clatot, F. (2014) CD70: A Potential Target in Breast Cancer? J Cancer. 5, 761–764

74. Kisor, K. P., Ruiz, D. G., Jacobson, M. P., and Barber, D. L. (2024) A role for pH dynamics regulating transcription factor DNA binding selectivity. 10.1101/2024.05.21.595212

75. Czowski, B. J., and White, K. A. (2024) Intracellular pH regulates β-catenin with low pHi increasing adhesion and signaling functions. 10.1101/2024.03.22.586349

76. Fang, Y., Liu, Z., Chen, Z., Xu, X., Xiao, M., Yu, Y., Zhang, Y., Zhang, X., Du, Y., Jiang, C., Zhao, Y., Wang, Y., Fan, B., Terheyden-Keighley, D., Liu, Y., Shi, L., Hui, Y., Zhang, X., Zhang, B., Feng, H., Ma, L., Zhang, Q., Jin, G., Yang, Y., Xiang, B., Liu, L., and Zhang, X. (2017) Smad5 acts as an intracellular pH messenger and maintains bioenergetic homeostasis. Cell Research. 27, 1083–1099

77. Cannell, I. G., Sawicka, K., Pearsall, I., Wild, S. A., Deighton, L., Pearsall, S. M., Lerda, G., Joud, F., Khan, S., Bruna, A., Simpson, K. L., Mulvey, C. M., Nugent, F., Qosaj, F., Bressan, D., CRUK IMAXT Grand Challenge Team, Dive, C., Caldas, C., and Hannon, G. J. (2023) FOXC2 promotes vasculogenic mimicry and resistance to anti-angiogenic therapy. Cell Rep. 42, 112791

78. Munro, D., Ghersi, D., and Singh, M. (2018) Two critical positions in zinc finger domains are heavily mutated in three human cancer types. PLoS Comput. Biol. 14, e1006290

79. Lund, L. M., Marchi, A. N., Alderfer, L., Hall, E., Hammer, J., Trull, K. J., Hanjaya-Putra, D., and White, K. A. (2024) Intracellular pH dynamics respond to microenvironment stiffening and mediate vasculogenic mimicry through β-catenin. 10.1101/2024.06.04.597454

80. Damaghi, M., Wojtkowiak, J. W., and Gillies, R. J. (2013) pH sensing and regulation in cancer. Front Physiol. 4, 370

81. Czowski, B. J., Romero-Moreno, R., Trull, K. J., and White, K. A. (2020) Cancer and pH Dynamics: Transcriptional Regulation, Proteostasis, and the Need for New Molecular Tools. Cancers (Basel). 12, E2760

82. Perez-Sala, D., Collado-Escobar, D., and Mollinedo, F. (1995) Intracellular alkalinization suppresses lovastatin-induced apoptosis in HL-60 cells through the inactivation of a pH-dependent endonuclease. J Biol Chem. 270, 6235–42

83. Matsumoto, A., Onoyama, I., Sunabori, T., Kageyama, R., Okano, H., and Nakayama, K. I. (2011) Fbxw7-dependent Degradation of Notch Is Required for Control of “Stemness” and Neuronal-Glial Differentiation in Neural Stem Cells. Journal of Biological Chemistry. 286, 13754–13764

84. Shao, S., Zhao, X., Zhang, X., Luo, M., Zuo, X., Huang, S., Wang, Y., Gu, S., and Zhao, X. (2015) Notch1 signaling regulates the epithelial–mesenchymal transition and invasion of breast cancer in a Slug-dependent manner. Mol Cancer. 14, 28

85. Bohin, N., Keeley, T. M., Carulli, A. J., Walker, E. M., Carlson, E. A., Gao, J., Aifantis, I., Siebel, C. W., Rajala, M. W., Myers, M. G., Jones, J. C., Brindley, C. D., Dempsey, P. J., and Samuelson, L. C. (2020) Rapid Crypt Cell Remodeling Regenerates the Intestinal Stem Cell Niche after Notch Inhibition. Stem Cell Reports. 15, 156–170

86. Clevers, H. (2013) The Intestinal Crypt, A Prototype Stem Cell Compartment. Cell. 154, 274–284

87. Yuan, X., Zhang, M., Wu, H., Xu, H., Han, N., Chu, Q., Yu, S., Chen, Y., and Wu, K. (2015) Expression of Notch1 Correlates with Breast Cancer Progression and Prognosis. PLoS ONE. 10, e0131689

88. Tsogtbaatar, E., Landin, C., Minter-Dykhouse, K., and Folmes, C. D. L. (2020) Energy Metabolism Regulates Stem Cell Pluripotency. Front. Cell Dev. Biol. 8, 87

89. Kang, H., Kim, H., Lee, S., Youn, H., and Youn, B. (2019) Role of Metabolic Reprogramming in Epithelial–Mesenchymal Transition (EMT). IJMS. 20, 2042

90. Bhattacharya, D., Azambuja, A. P., and Simoes-Costa, M. (2020) Metabolic Reprogramming Promotes Neural Crest Migration via Yap/Tead Signaling. Developmental Cell. 53, 199–211.e6

91. Mosier, J. A., Schwager, S. C., Boyajian, D. A., and Reinhart-King, C. A. (2021) Cancer cell metabolic plasticity in migration and metastasis. Clin Exp Metastasis. 38, 343–359

92. Druzhkova, I. N., Shirmanova, M. V., Lukina, M. M., Dudenkova, V. V., Mishina, N. M., and Zagaynova, E. V. (2016) The metabolic interaction of cancer cells and fibroblasts – coupling between NAD(P)H and FAD, intracellular pH and hydrogen peroxide. Cell Cycle. 15, 1257– 1266

93. Hanahan, D., and Weinberg, R. A. (2011) Hallmarks of cancer: the next generation. Cell. 144, 646–74

94. Liberti, M. V., and Locasale, J. W. (2016) The Warburg Effect: How Does it Benefit Cancer Cells? Trends in Biochemical Sciences. 41, 211–218

95. Zhou, W., Choi, M., Margineantu, D., Margaretha, L., Hesson, J., Cavanaugh, C., Blau, C. A., Horwitz, M. S., Hockenbery, D., Ware, C., and Ruohola-Baker, H. (2012) HIF1α induced switch from bivalent to exclusively glycolytic metabolism during ESC-to-EpiSC/hESC transition: Metabolic switch in ESC-to-EpiSC/hESC transition. The EMBO Journal. 31, 2103– 2116

